# Collagen mineralization decreases NK cell-mediated cytotoxicity of breast cancer cells via increased glycocalyx thickness

**DOI:** 10.1101/2024.01.20.576377

**Authors:** Sangwoo Park, Siyoung Choi, Adrian A. Shimpi, Lara A. Estroff, Claudia Fischbach, Matthew J. Paszek

## Abstract

Skeletal metastasis is common in patients with advanced breast cancer, and often caused by immune evasion of disseminated tumor cells (DTCs). In the skeleton, tumor cells not only disseminate to the bone marrow, but also to osteogenic niches in which they interact with newly mineralizing bone extracellular matrix (ECM). However, it remains unclear how mineralization of collagen type I, the primary component of bone ECM, regulates tumor-immune cell interactions. Here, we have utilized a combination of synthetic bone matrix models with controlled mineral content, nanoscale optical imaging, and flow cytometry to evaluate how collagen type I mineralization affects the biochemical and biophysical properties of the tumor cell glycocalyx, a dense layer of glycosylated proteins and lipids decorating their cell surface. Our results suggest that collagen mineralization upregulates mucin-type O-glycosylation and sialylation by tumor cells, which increased their glycocalyx thickness while enhancing resistance to attack by Natural Killer (NK) cells. These changes were functionally linked as treatment with a sialylation inhibitor decreased mineralization-dependent glycocalyx thickness and made tumor cells more susceptible to NK cell attack. Together, our results suggest that interference with glycocalyx sialylation may represent a therapeutic strategy to enhance cancer immunotherapies targeting bone-metastatic breast cancer.

## 1. Introduction

Metastasis is the leading cause of death in patients with advanced breast cancer and develops in the skeleton in over 70% of these patients[1]. Because clinically evident bone metastasis is challenging to treat, better understanding the mechanisms that regulate the fate of disseminated tumor cells (DTCs) prior to lesion formation promises to inform therapeutic options to prevent metastasis. Tumor cells spread to bone early during disease and often remain quiescent for years, decades, or sometimes forever[2]. The signals that control whether or not quiescent tumor cells proceed to form metastasis remain poorly understood but altered immunoregulatory mechanisms play a key role in this process[3]. Indeed, latent tumor cells display increased propensity to evade immune cell-mediated cytotoxicity. For example, bone-resident dormant DTCs in breast cancer patients without detectable metastases are lined by higher number of immune cells[4], and latent metastatic cancer cells evade clearance by the immune system[5]. While prior work identified how tumor cell intrinsic signals regulate interactions with immune cells, it is less well understood how bone extracellular matrix (ECM) controls immune surveillance of early DTCs. Clarifying these connections is important as disseminated bone-metastatic tumor cells often target osteogenic niches in which they interact with bone matrix and because bone matrix properties can impact tumor cell latency[6,7].

Bone matrix is unique in its physicochemical properties, and ECM physicochemical properties can control tumor-immune cell interactions[8–10]. Therefore, bone ECM may be implicated in the immunoregulation of skeletal metastasis. Studying these possible connections requires biomaterial models that mimic relevant bone matrix properties for *in vitro* studies. During osteogenesis, bone forming cells first secrete osteoid, a fibrillar collagen type I-rich organic matrix into which carbonated hydroxyapatite (HAp; Ca_10_(PO_4_CO_3_)_6_(OH)_2_) nanoparticles are subsequently deposited via both intrafibrillar and extrafibrillar mineralization[11,12]. Several strategies have been developed to mimic the materials properties of mineralized collagen, but many of these approaches deposit HAp on top of synthetic biomaterials or use protocols that result in extrafibrillar rather than intrafibrillar collagen mineralization, which is a primary mechanism during bone formation[13]. The polymer-induced liquid precursor (PILP) method can mimic physiological intrafibrillar collagen mineralization using polyelectrolytes such as polyaspartic acid (PAA) to simulate the function of acidic proteins *in vivo*[14,15]. We have previously adapted the PILP method to develop cell culture substrates for studies of bone metastasis and identified that collagen mineralization inhibits tumor growth by increasing tumor cell stemness, a phenotype that is immune privileged[7,16], but how these changes affect immune surveillance remains unclear.

Cancer cells are encased in a dense layer of glycosylated proteins and lipids called the glycocalyx, which plays a powerful role in the establishment of a permissive tumor immune microenvironment (TIME)[17–20]. Essentially all cancers exhibit changes in glycan synthesis, resulting in glycans at atypical levels or with altered structural attributes[21–23]. Relatively small changes in the expression of even a few glycosyltransferases that govern glycosylation events can have substantial consequences in cancer. Among the immunomodulatory glycans, sialic acid containing glycans (sialoglycans) on tumor cells can function as potent suppressors of effector cells in the TIME[24,25]. For instance, tumor sialoglycans can ligate the sialic acid-binding immunoglobulin-like lectin (Siglec) family of immunomodulatory receptors that are expressed in most immune cell types[26,27]. Aberrantly high expression of sialoglycans and other cancer-associated glycosylation patterns are implicated in multiple stages of bone metastasis, including extravasation from the bone vasculature, early-stage colonization in the bone, and later outgrowth[28,29]. However, whether the bone microenvironment’s unique features, such as the presence of mineralized matrix, regulates glycosyltransferase expression, glycan biosynthesis, or the construction of a cancer-cell glycocalyx with altered immune-related functionality has not been directly tested.

Beyond serving as immunomodulatory ligands, glycans contribute to the steric, electrostatic, and hydrophilic interactions that shape the swollen architecture of the glycocalyx layer[30]. Effector immune cells engage tumor cells through tight, receptor-mediated interactions that are obligatory for cell-mediated cytotoxicity. A thick glycocalyx is proposed to function as a barrier that shields tumor cell ligands from engagement by the receptors of surveilling immune cells[31]. Typical immune receptors extend from the effector cell surface by 10-20 nm, whereas the thickness of the cancer cell glycocalyx can exceed 100 nm[32–34]. Theoretical models of cell adhesion have long predicted a potentially powerful role for the glycocalyx layer in regulating specific interactions between cells mediated by receptors of a finite size, with the nanometer-scale thickness of the glycocalyx emerging as a key physical parameter[35,36]. Cancer aggression and poor patient outcomes correlate with the expression of large glycoproteins that can contribute to abnormal glycocalyx thickening, which is implicated in altered cell adhesion, migration, invasiveness, growth, and susceptibility to DNA damage[33,34,37–39]. Notably, solid tumor cells are often densely grafted with cell-surface mucins, a family of membrane-anchored biopolymers that are defined by their extensive O-glycosylation[40]. When overexpressed, cell-surface mucins and the glycans that they carry serve as major structural elements in the cancer-cell glycocalyx, contributing to its aberrant thickening[33,41]. Multiple studies have implicated mucin expression and changing glycosylation patterns in protection of tumor cells from NK cells and cytotoxic T lymphocytes[17,42,43]. However, whether specific bone matrix properties, such as mineralization, can instruct tumor cells to modulate their glycocalyx structure in a manner that supports immune evasion remains poorly understood due in part to a lack of integrated approaches that enable recapitulation of relevant bone matrix properties and analysis of its effects on the nanoscale glycocalyx structure.

Here, we used a combination of synthetic bone matrix models with controlled mineral content, nanoscale optical imaging, and selective manipulation of tumor cell glycan patterns to define how collagen mineralization regulates the breast cancer cell glycocalyx and to determine which functional consequences these changes have on NK cell cytotoxicity. More specifically, we cultured various breast cancer cell lines on collagen and mineralized collagen to determine their glycocalyx-related phenotypic and gene expression changes, which we related to changes in glycocalyx composition and thickness using flow cytometry and Scanning Angle Interference Microscopy (SAIM)[44,45], respectively. Using both human primary NK cells and the NK-92 cell line, we subsequently determined how the detected changes affect NK cell-mediated cytotoxicity and validated the relevance of our conclusions using glycocalyx-editing enzymes and inhibitors. Our results indicate that collagen mineralization alters glycan expression to increase glycocalyx thickness and that these changes, in turn, increase resistance to NK cell attack. Collectively, our findings suggest bone ECM plays a critical role in modulating immune response during early stages of metastasis with potential implications for how tumor cell latency in the skeleton may be targeted therapeutically.

## 2. Results

### 2.1. Bone metastatic potential and glycocalyx thickness of breast cancer cells are inversely related to NK cell-mediated cytotoxicity

Although altered glycosylation is a hallmark of tumor cells and associated with decreased immune response[46,47], it remains unclear if the glycocalyx of tumor cells varies as a function of their bone-metastatic potential and how these changes affect their interactions with immune cells. We investigated these connections with three breast cancer cell lines that differ in their ability to form skeletal lesions in mice. MCF7 are estrogen-receptor positive breast cancer cells, the major breast cancer subtype prone to develop bone metastasis in patients, but these cells are non-metastatic and express genes associated with dormancy[48,49]. In contrast, triple negative MDA-MB231 cells are widely used to model bone metastasis in mice via intracardiac injection. The highly bone metastatic cell line BoM-1833 was established from bone metastases that formed in nude mice following inoculation with the MDA-MB231 cell line.[50]. To measure glycocalyx thickness of these different breast cancer cell lines, we subjected them to SAIM, which maps the thickness of a cell’s glycocalyx with 5-10 nm precision by imaging and quantifying the distance between two fluorescent molecules of which one is adsorbed to a reflective cell substrate and the other is labeling the cell’s membrane, the gap between the two corresponding to the glycocalyx thickness[41,44,51] (Fig. 1a). Interestingly, glycocalyx thickness was highest for BoM-1833 (77 nm) and decreased for MDA-MB231 (68 nm) and MCF7 (59 nm) (Fig. 1b) suggesting that glycocalyx thickness positively correlates with the bone metastatic potential of these different cell lines.

**Figure 1.**
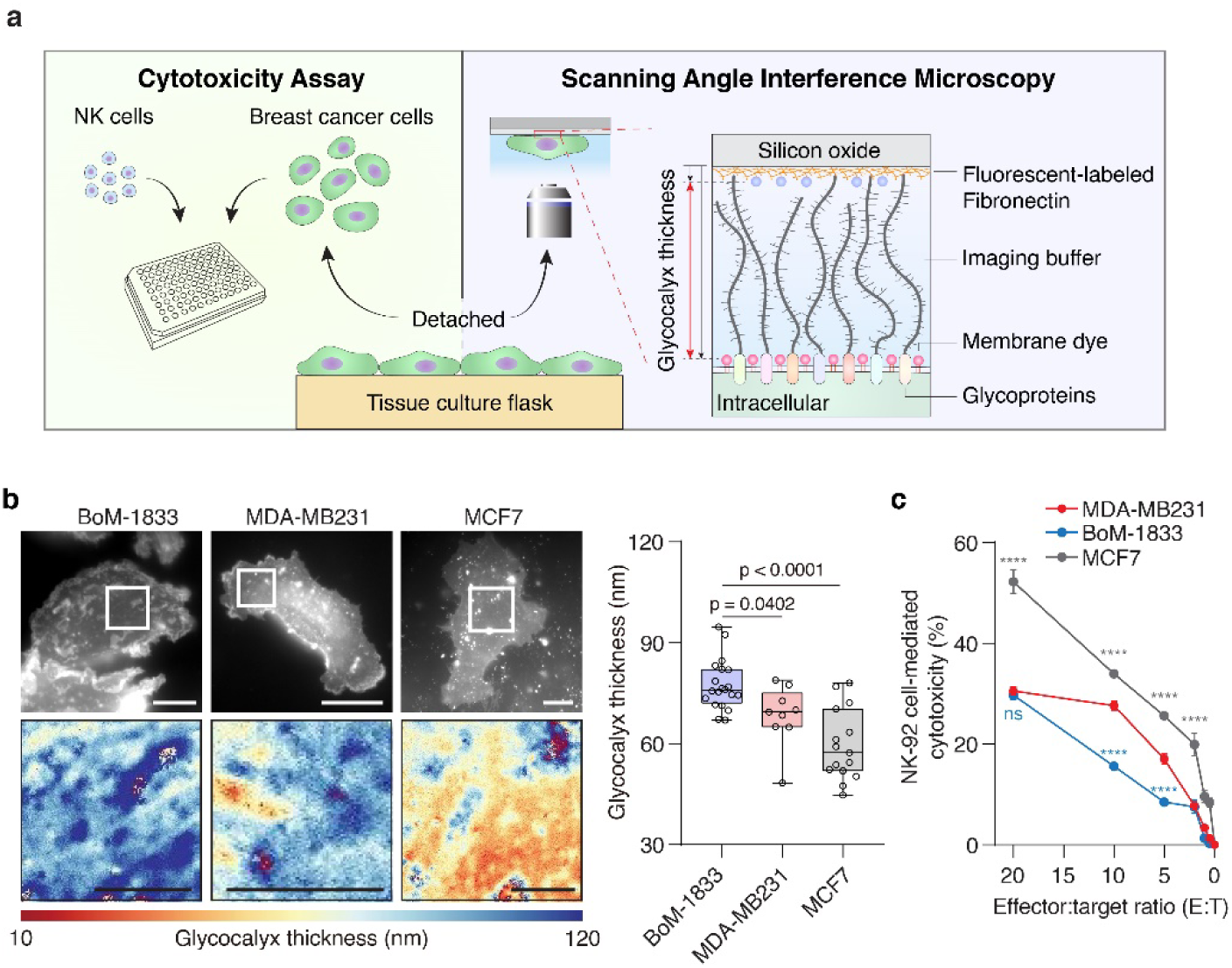
Glycocalyx thickness and NK-92 cell-mediated cytotoxicity of breast cancer cell lines. (a) Schematic of experimental setup to measure glycocalyx thickness using Scanning Angle Interference Microscopy (SAIM) and NK cell-mediated cytotoxicity. (b) Representative widefield images and glycocalyx thickness map reconstructed by SAIM (left) and analysis (right) of breast cancer cell lines precultured on polystyrene (PS). Scale bars, 10 µm (top) and 5 µm (bottom). Boxes and whiskers show the first and third quartiles (boxes), median, and range of the data. Each condition includes a minimum of 9 cells from a representative experiment. Statistical analysis by one-way ANOVA with Tukey’s post-hoc tests. (c) NK-92 cell-mediated cytotoxicity of breast cancer cell lines at the indicated effector cell and target cell (E:T) ratio; an E:T ratio of 0 indicates that NK-92 cells-mediated cytotoxicity were assayed in the absence of NK-92 cells. Results are the mean ± s.d. (*n* = 3). Statistical analysis by one-way ANOVA with Tukey’s post-hoc tests. Statistical analysis under E:T ratio 2 is not shown. Statistical significance is given by **** p < 0.0001.

Because we have previously identified that cells with a thicker glycocalyx can evade immune cell-mediated cytotoxicity more effectively than their counterparts with a less dense glycocalyx[41], we hypothesized that BoM-1833 may resist immune attack more effectively than MDA-MB231 and even more so than MCF7. We tested this hypothesis with NK cells because they play key roles in the innate immune response, control metastasis formation, and are increasingly explored for adoptive cell therapy[52,53]. In addition, it has been shown that tumors can develop mechanisms to resist attack by NK and (Chimeric Antigen Receptor) CAR-NK cells, and changes in glycocalyx structure may contribute to this process[41]. To examine NK cell-mediated cytotoxicity, we incubated the human NK-92 cell line with the different tumor cell lines at various effector (E; NK-cells) to target (T; tumor cells) ratios (Fig. 1c). Regardless of E:T ratio, MCF7 cells were killed by NK cells much more effectively than MDA-MB231 and BoM-1833 cells. At high and very low E:T ratios, NK cells killed MDA-MB231 and BoM-1833 cells at similar levels. However, at intermediate E:T ratios BoM-1833 resisted NK cell attack more efficiently than MDA-MB231. Collectively, these data suggest breast cancer cells that develop bone metastasis in mouse models have increased glycocalyx thickness and are more resistant to NK cell-mediated cytotoxicity relative to cells that disseminate to bone, but do not reliably form metastatic lesions.

### 2.2. Collagen mineralization alters MDA-MB231 immunoregulation and glycocalyx formation

During early stages of bone metastasis, breast cancer cells often target osteogenic niches in which they interact with mineralizing matrix[6]. We have previously identified that interactions with bone matrix affect tumor cell phenotype differently than culture on plastic surfaces[7], and others have shown that cell-ECM interactions regulate glycocalyx formation[17–19,54]. Given these connections, we speculated that bone matrix mineralization may regulate aspects of glycocalyx formation and immunoregulation. To examine this possibility, we used collagen substrates with and without intrafibrillar mineralization to mimic physiologic bone matrix mineralization and scenarios in which newly deposited bone matrix is not properly mineralized, respectively (e.g. due to aging or chemotherapy)[55–57]. We fabricated these substrates with the PILP method, using our previously established protocols[14]. With this method, collagen fibers are covalently attached to glass coverslips mounted onto a polydimethylsiloxane (PDMS) base and then submerged in the mineral-forming solution in an upside-down configuration to ensure intrafibrillar mineralization rather than mineral precipitation on top of collagen fibers (Fig. 2a). Intrafibrillar mineralization was initiated with a solution containing calcium, phosphate, and PAA to form nano-scale amorphous calcium phosphate that infiltrates the collagen fibrils and subsequently matures to HAp as we have confirmed previously using Fourier transform infrared spectroscopy and X-ray diffraction [7,58]. Scanning electron microscopy (SEM) and backscattered electron (BSE) image analysis further confirmed that our protocols led to the formation of thicker, mineral-containing collagen fibers without altering the overall architecture of the collagen network (Fig. 2b).

**Figure 2.**
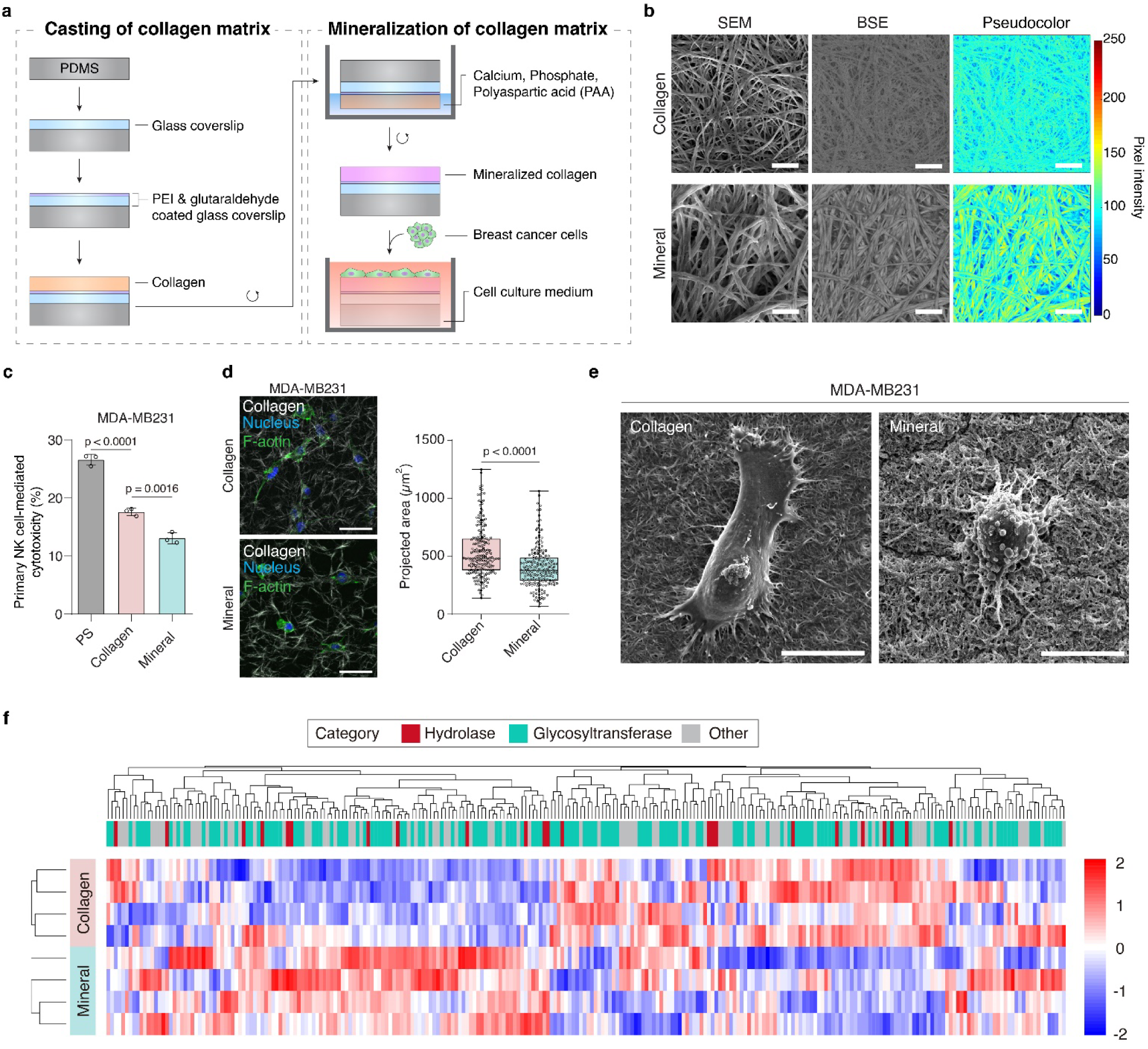
Engineered bone-matrix model to study the effect of collagen mineralization on phenotypic and gene expression changes of MDA-MB231. (a) Schematic showing the fabrication of mineralized collagen type I matrices using the polymer-induced liquid-precursor (PILP) method. (b) Representative scanning electron microscopy (SEM) and backscatter electron (BSE) images showing mineralization-dependent changes of collagen fiber structure and mineral content, respectively. Pseudocolor indicates mineral content as determined from BSE images by converting grey-scale pixel intensity to a linear 256-bit colour scale. Scale bars, 2 µm. (c) Primary NK cell-mediated cytotoxicity of MDA-MB231 cells precultured on PS, collagen, and mineralized collagen (mineral) at a 10:1 E:T ratio. Results are the mean ± s.d. (*n* = 3). (d) Representative confocal images and corresponding quantification of cell morphology by image analysis. Scale bars, 50 µm. Boxes and whiskers show the first and third quartiles (boxes), median, and range of the data. Each condition includes a minimum of 202 cells from a representative experiment. (e) SEM images of MDA-MB231 cells cultured on collagen or mineralized collagen. Scale bars, 10 µm. (f) Heatmap of glycogenes (categorized as glycosyltransferase, hydrolase, and other) differentially expressed by MDA-MB231 cells on collagen versus mineralized collagen. GEO accession number is GSE229094. Results are the mean ± s.d. (*n* = 3). In panel **c** and **d**, statistical differences were determined using one-way ANOVA with multiplicity-adjusted p values from Tukey’s multiple comparisons test (c) and two-tailed unpaired t-test with Welch’s correction (d).

To test how collagen mineralization affects the ability of breast cancer cells to resist NK-cell mediated cytotoxicity, we cultured MDA-MB231 breast cancer cells on mineralized and control collagen. Subsequently, cells were harvested from these different substrates and subjected to a NK cell cytotoxicity assay. Here, we used primary NK cells isolated from human blood to ensure patient relevance and incubated these cells with tumor cells at a E:T ratio of 10 informed by our prior experiments with NK-92 cells (Fig. 1c). MDA-MB231 cells precultured on mineralized collagen were more resistant to NK cell attack than cells precultured on non-mineralized collagen or polystyrene, highlighting the importance of relevant cell culture substrates when studying immune cell responses. (Fig. 2c). Although various mechanisms could be responsible for these findings (e.g. including lytic granules-mediated apoptosis or death receptor-mediated cytotoxicity), substrate-dependent changes of cell morphology suggested that altered glycocalyx formation could also play a role. More specifically, MDA-MB231 cells on mineralized collagen were smaller and rounder relative to cells on collagen (Fig. 2d) consistent with previous reports that increased glycocalyx density leads to decreased cell spreading[33,59]. Moreover, SEM imaging revealed that MDA-MB231 cells on mineralized collagen displayed prominent membrane protrusions (Fig. 2e), which we had previously shown can result from glycocalyx-dependent membrane bending[60].

To more directly determine if interactions of MDA-MB231 cells with mineralized versus control collagen alter glycocalyx formation, we took advantage of our previously published RNA-seq dataset that describes the transcriptome of MDA-MB231 cells cultured on mineralized versus control collagen[7]. Bioinformatic analysis of this dataset followed by hierarchical clustering suggested that glycogenes involved in the synthesis and breakdown of glycans and categorized as glycosyltransferases, hydrolases, and other glycan related genes were expressed differentially between MDA-MB231 cells on mineralized collagen relative to non-mineralized collagen (Fig. 2f; Supplementary Fig. 1). Collectively, these results suggest that MDA-MB231 cells interacting with mineralized versus non-mineralized collagen are more resistant to NK cell attack and exhibit morphological and gene expression differences suggestive of glycocalyx changes.

### 2.3. Collagen mineralization increases tumor cell glycocalyx thickness and evasion of NK cell-mediated cytotoxicity

Given that MDA-MB231 cells cultured on mineralized collagen substrates evaded NK cell-mediated killing more effectively than their counterparts on non-mineralized collagen and that these cells displayed morphological and gene expression changes indicative of glycocalyx changes, we next tested how collagen mineralization affects glycocalyx thickness and whether these measurements correlated with NK cell-mediated cytotoxicity. Consistent with the determined gene expression and morphological changes, SAIM revealed that the glycocalyx of MDA-MB231 cells cultured on mineralized collagen was significantly thicker (69 nm) than the glycocalyx of the same cells cultured on collagen (50 nm) (Fig. 3a). Given that increased glycocalyx thickness can promote metastatic potential by increasing cell survival under adhesion-limited conditions[33,61], and because tumor cells interacting with mineralized collagen adhere less strongly[7,58], these results suggest that collagen mineralization promotes cell survival and metastatic potential by increasing glycocalyx thickness. In addition to changing these tumor cell intrinsic features, we found that MDA-MB231 cells precultured on mineralized collagen were also more resistant to NK-92 cell-mediated cytotoxicity (34.9%) than their counterparts cultured on collagen (42.4%) (Fig. 3b). Hence, our data indicate that changes in bone matrix mineralization may regulate metastasis formation by controlling multiple hallmarks of the TIME synergistically.

**Figure 3.**
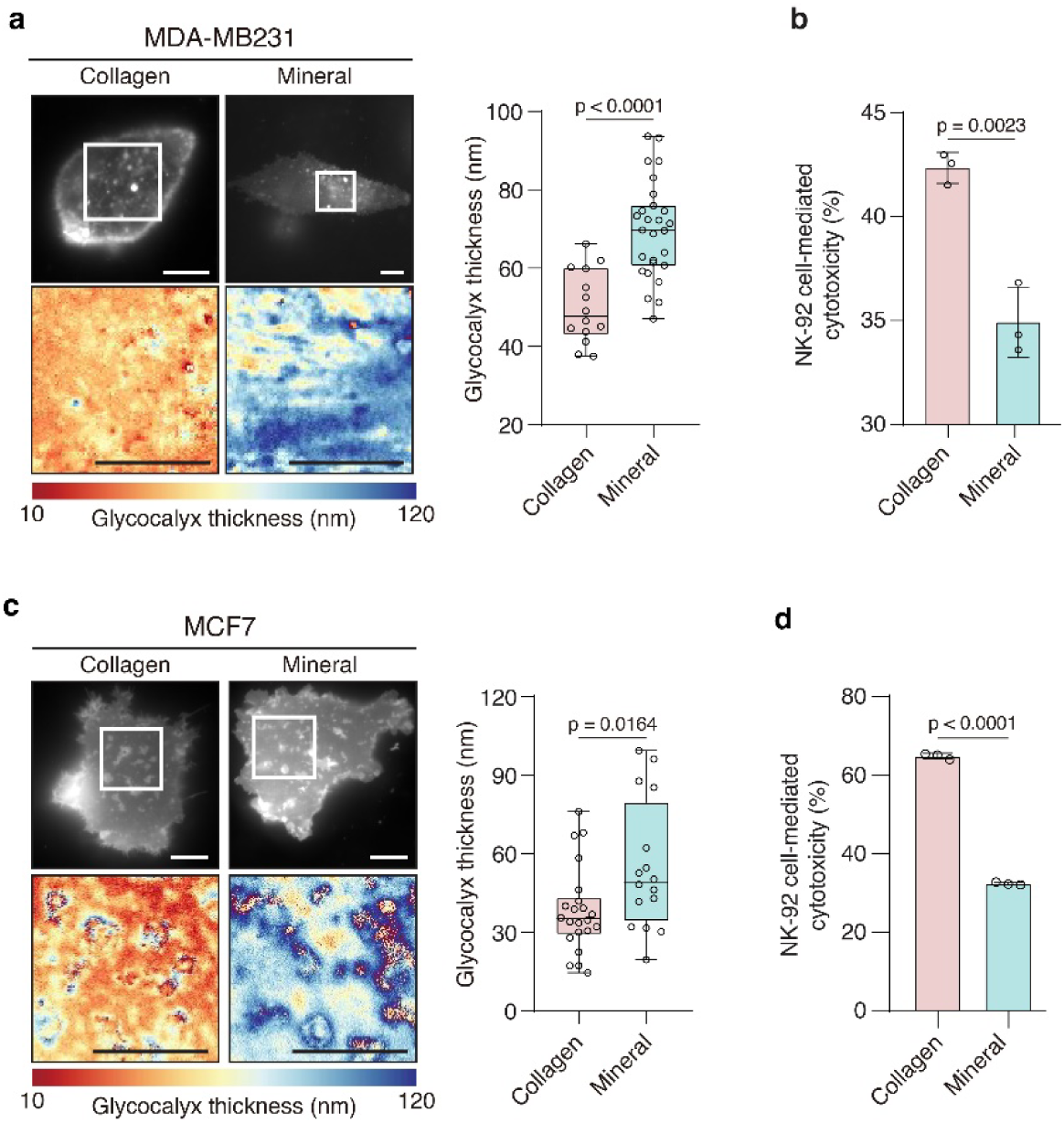
Effects of collagen mineralization on glycocalyx thickness and protection against NK-92 cell-mediated cytotoxicity in MDA-MB231 and MCF7 cells. (a) Representative widefield images and glycocalyx thickness maps reconstructed by SAIM (left) and corresponding analysis (right) of MDA-MB231 cells precultured on collagen and mineralized collagen. Scale bars, 10 µm. (b) NK-92 cell-mediated cytotoxicity of MDA-MB231 cells at a E:T ratio of 10:1. (c) Representative widefield images and glycocalyx thickness maps reconstructed by SAIM (left) and corresponding analysis (right) of MCF7 precultured on collagen and mineralized collagen. Scale bars, 10 µm. (d) NK-92 cell-mediated cytotoxicity of MCF7 at a E:T ratio of 5:1. In panel **a** and **c**, boxes and whiskers show the first and third quartiles (boxes), median, and range of the data. Each condition includes a minimum of 14 cells from a representative experiment. In panel **b** and **d**, results are the mean ± s.d. (*n* = 3). In panel **a**-**d**, statistical differences were determined using two-tailed unpaired t-test.

To test whether our findings were limited to MDA-MB231 cells or more broadly relevant, we repeated these experiments with BoM-1833 and MCF7 cells (Fig. 3 and Supplementary Fig. 2). Compared to the parental MDA-MB231 cell line, the bone-metastatic BoM-1833 line exhibited a thicker glycocalyx on both collagen (79 nm) and mineralized collagen (83 nm), consistent with their greater resistance in general to NK-cell mediated cytotoxicity (Fig. 1c and Supplementary Fig. 2). We next investigated MCF7 as an ER+ breast cancer model that can disseminate to bone but is known to often assume dormancy under these conditions[62]. We found previously that interactions with mineralized collagen induce a more stem cell-like phenotype in MCF7 cells[7], a phenotype that has been correlated by others with changes in glycocalyx formation[63]. While MCF7 cells cultured on mineralized collagen had a 55 nm thick glycocalyx, the same cells cultured on collagen had a 38 nm thick glycocalyx (Fig. 3c). Given the relatively thin glycocalyx structure of MCF7 cells, we speculated that these cells may be more susceptible to attack by NK-92 cells compared to MDA-MB231. Indeed, at the 10:1 E:T ratio used for experiments with MDA-MB231 most MCF7 cells died (data not shown). Hence, we performed the cytotoxicity assay at a significantly reduced E:T ratio (5:1). Similar to the result of MDA-MB231, we found that MCF7 precultured on mineralized collagen were also more protected against NK-92 cell-mediated cytotoxicity compared to the same cells in the collagen condition (Fig. 3d). Together, these results suggest that collagen mineralization increases glycocalyx thickness and resistance to NK-92 cell-mediated killing in both MDA-MB231 and MCF7 cells, but that MCF7 are much more susceptible to NK cell attack suggesting that changes in glycocalyx thickness scale with NK cell response.

### 2.4. Mucins and sialoglycans protect cancer cells on mineralized collagen against effector immune cells

We next sought to unravel the molecular changes underlying the thicker glycocalyx and immune protection that was observed in the previous experiments. We noted multiple, significant changes in the expression of genes that could potentially regulate the glycocalyx structure in mineralized conditions (Supplementary Fig. 1). Glycosylation machinery that was significantly upregulated included α2,6 sialyltransferases for O-glycans (ST6GALNAC1, p = 0.439, and ST6GALNAC2, p = 0.0009), governing enzymes for LacDiNac extension of O-and N-glycans (B4GALNT3, p < 0.0001 and B4GALNT4, p = 0.0019), and enzymes for hyaluronic acid (HA) anabolism (HAS2, p < 0.0001) and catabolism (HYAL2, p = 0.0042). We also observed significant upregulation of some mucin genes, including cancer-associated MUC1 (p < 0.0001), and members of the galectin family of multivalent lectins that are implicated in crosslinking mucins and other glycoproteins (LGALS1, p < 0.0001, and LGALS3, p < 0.0001). These results suggested that the observed structural changes of the glycocalyx in mineralized conditions could be linked to one or more potential candidates: HA biosynthesis, sialylation, N-glycosylation, and O-glycosylated mucins.

To dissect the contribution of each component to the glycocalyx structure and its immune protective effects, we used enzymatic tools to selectively remove cell-surface HA, sialic acids, N-glycans, and O-glycosylated mucins from cells cultured on the different bone matrix models (Fig. 4a). We initially tested the contributions of these glycocalyx elements in the protection against cell-mediated cytotoxicity by effector cells. Similar to the experiments described above, NK-92 were selected as the effector cell model because they generally lack glycan-binding inhibitory receptors, like Siglec-7 and –9, which might otherwise inhibit the immune cells directly in response to ligation by tumor cell glycans[42,64,65]. In co-culture assays of NK-92 cells with MDA-MB231 target cells precultured on the different collagen substrates, we observed no significant changes in cytotoxicity when the N-glycans were selectively removed from the target cells by PNGase F treatment or when HA was removed with hyaluronidase (Fig. 4b and Supplementary Fig. 3, 5). However, we did observe significant increases in NK-92 cell-mediated cytotoxicity against the cancer cells when their O-glycosylated mucins were trimmed with the StcE mucinase or when treated with the *Salmonella typhimurium* sialidase, which removes sialic acids with α2,3, α2,6, and α2,8 linkages (Fig. 4b,c). Given the nonsignificant effects of N-glycan and HA removal, these results implicated mucins and mucin sialoglycans in collagen mineralization-dependent immune evasion.

**Figure 4.**
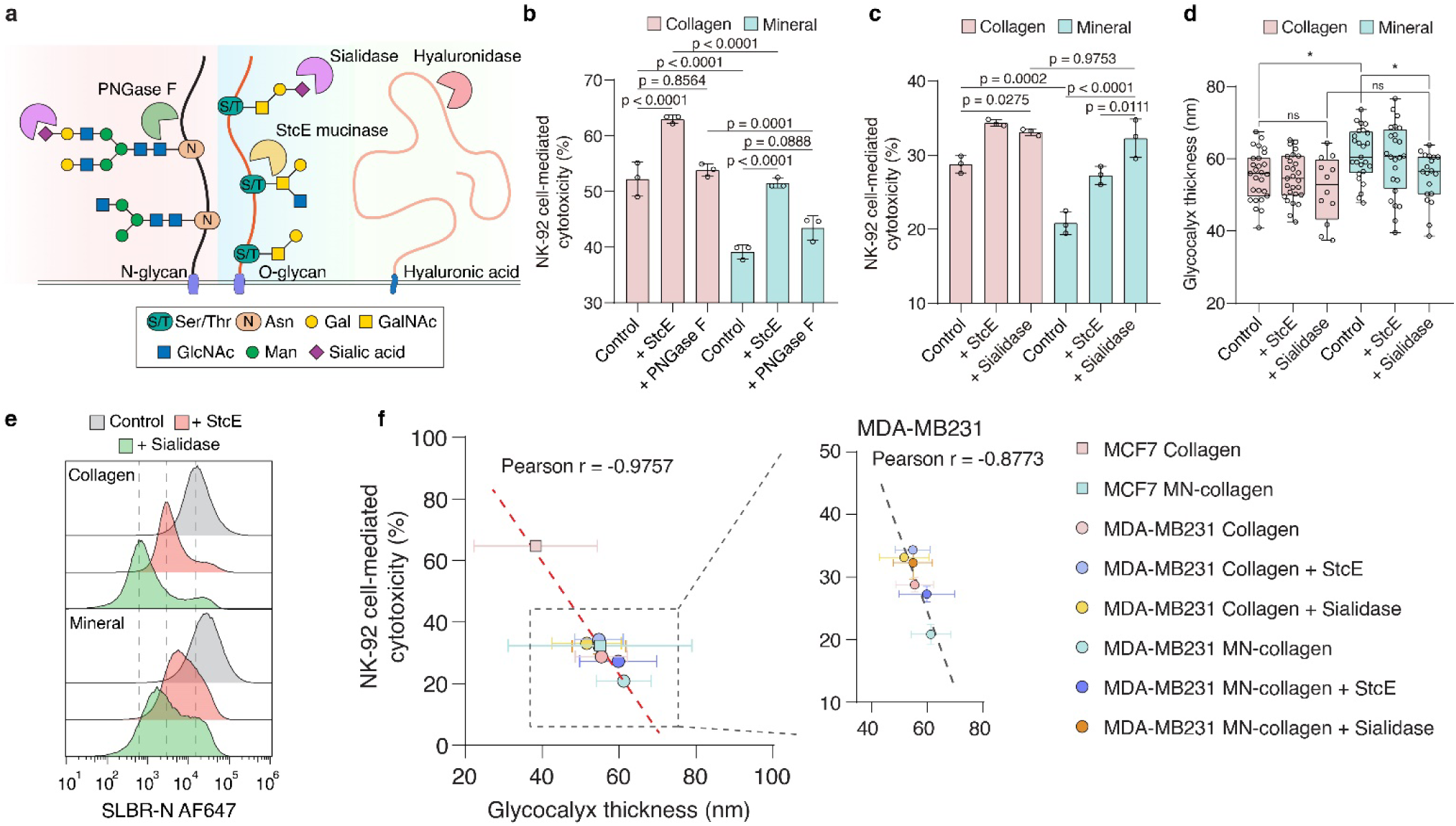
Enzymatic removal of cell-surface mucins and mucin sialoglycans on collagen mineralization enhances NK cell cytolytic efficiency via reduced glycocalyx thickness. (a) Cartoon showing the enzyme-mediated cleavage of different glycan structures. (b) NK-92 cell-mediated cytotoxicity against MDA-MB231 cells treated with control buffer or 100 nM StcE, or 5,000 U/mL PNGase F prior to co-culture. E:T ratio is 10:1. (c) NK-92 cell-mediated cytotoxicity against MDA-MB231 cells treated with control buffer or 100 nM StcE, or 100 nM sialidase prior to co-culture. E:T ratio is 10:1. (d) Quantification of mean glycocalyx thickness in MDA-MB231 cells before and after treatment with 100 nM StcE or 100 nM sialidase. Boxes and whiskers show the first and third quartiles (boxes), median, and range of the data. Each condition includes a minimum of 12 cells from a representative experiment. (e) Flow cytometry analysis of SLBR-N level on MDA-MB231 cells after treatment with 100 nM StcE and 100 nM sialidase. (f) Inverse correlation between NK-92 cell-mediated cytotoxicity and glycocalyx thickness from Fig. 3c,d and Fig. 4b-d with Pearson correlation coefficient, r = –0.9757. Dashed line indicates a linear fit with the data. In panel **b** and **c**, results are the mean ± s.d. (*n* = 3). In panel **b**-**d**, statistical differences were determined using one-way ANOVA with multiplicity-adjusted p values from Tukey’s multiple comparisons test (b,c) and from Newman-Keuls’ multiple comparisons test (d).

We next tested whether O-glycosylated mucins and sialoglycans contributed to the fortification of the glycocalyx layer that was observed in mineralized conditions. We measured the glycocalyx thickness of cells cultured on non-mineralized or mineralized collagen following treatment with the panel of glycocalyx-editing enzymes (Fig. 4d). Sialidase treatment of cells cultured on mineralized collagen reduced their glycocalyx thickness by 6.4 nm compared to non-treated cells grown under similar conditions (Fig. 4d). Treatment with the StcE mucinase did not affect glycocalyx thickness in cells cultured on non-mineralized versus mineralized collagen substrates. However, StcE may have removed only a subset of glycoproteins displaying mucin-type O-glycans from the cell surface, consistent with reports that StcE does not cleave all mucin-domain containing glycoproteins[66,67]. Probing the StcE-treated cells with SLBR-N lectin, a mucin probe that binds preferentially to sialylated Core-II mucin-type O-glycans and has been used as a marker for cancer stem cells[63], we observed only a partial loss of probe reactivity to the cell surface following enzyme treatment (Fig. 4e and Supplementary Fig. 4a). A more pronounced loss in SLBR-N reactivity was observed following treatment of the cancer cells with sialidase, confirming removal of sialic acids from the Core-II O-glycans (Fig. 4e and Supplementary Fig. 4a). While the broad-spectrum *S. typhimurium* sialidase was expected to also remove the sialic acids of N-glycans, the lack of change in cytotoxicity against target cells treated with PNGase F suggested that sialylated N-glycans were not a major determinant of the protective function of the glycocalyx. Likewise, trimming of sialic acids from glycolipids, like gangliosides, by the sialidase would not be expected to significantly alter the extended glycocalyx thickness, since glycolipids extend only a couple nanometers from the cell surface, whereas the measured glycocalyx thickness of the cancer cells ranged from approximately 40-80 nm depending on the culture conditions and treatments. Taken together, these results implicated the importance of sialylated mucins to the glycocalyx structure and immune protection functions.

We considered to what extent the changes in glycocalyx thickness across the different conditions could explain the differences in the cancer cell susceptibility to effector cell killing. Compiling the data from Figure 3c,d and Figure 4b-d, we constructed a plot of NK-92 cell-mediated cytotoxicity versus the measured glycocalyx thickness of the cancer cells cultured on the various ECM substrates and subjected to different enzymatic treatments. We identified an inverse correlation between cytotoxicity and glycocalyx material thickness across the combined dataset spanning the range of conditions with a Pearson coefficient of –0.98 (Fig. 4f). The statistical outcome, a correlation coefficient near negative unity, indicated that the glycocalyx thickness parameter was strongly predictive of the cellular susceptibility to NK cell-mediated cytotoxicity. The strong statistical correlation between glycocalyx thickness and cytotoxicity implicated collagen mineralization in immune evasion by informing cancer cells to construct a glycocalyx with altered physical properties.

### 2.5. Collagen mineralization regulates sialoglycans and mucin O-glycans

Given the importance of mucins and mucin sialoglycans to the immune protective functions of the glycocalyx, we investigated how the composition of cell-surface O-glycans changes in response to cellular interactions with mineralized collagen. Following MDA-MB231 culture on mineralized and non-mineralized collagen substrates, mucin-type O-glycans and sialoglycans were probed with a panel of lectins and quantified by flow cytometry and confocal microscopy (Fig. 5a). In mineralized conditions, we observed substantial increases in the mucin-type core-I O-glycans, which were probed with peanut agglutinin (PNA). The core-1 O-glycans were highly sialylated, as indicated by the strong increase in cellular reactivity to PNA following sialidase treatment (Fig. 5b). Likewise, cells cultured on mineralized collagen showed a small increase in total cell-surface sialylation (α2,3, α2,6, and α2,8 linkages), which was measured using the periodate oxidation and aniline-catalyzed oxime ligation (PAL) method[68]. This strategy introduced aldehydes into sialic acids, enabling the labeled glycans to be subsequently probed with aminoxy-functionalized fluorophores (Fig. 5c).

**Figure 5.**
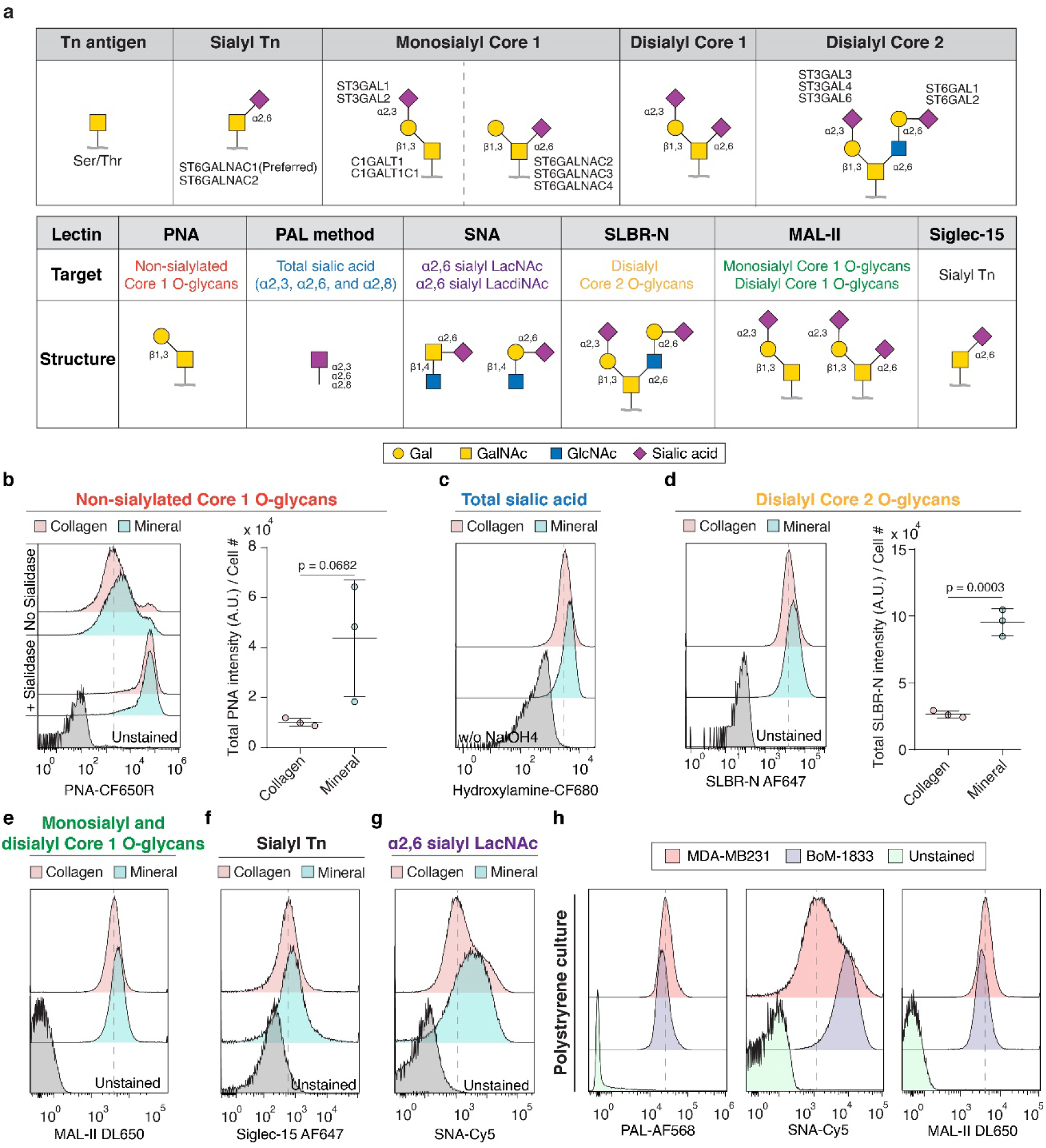
Collagen mineralization enhances the expression of sialoglycans and mucin O-glycans in metastatic breast cancer cells. (a) Schematic of representative glycosyltransferase genes for Tn-antigen, Sialyl Tn, monosialyl Core 1, disialyl Core 1, and disialyl Core 2 (top) and lectin probes targeting different glycan structures (bottom). (b) Flow cytometry histograms and quantification of PNA levels on MDA-MB231 cells treated with control buffer or 100 nM sialidase for 1 hour; PNA levels quantified by measurement of mean fluorescent intensity in confocal microscopy images. (c) Flow cytometry analysis of total cell-surface sialic acids on MDA-MB231 cells using the periodate oxidation and aniline-catalyzed oxime ligation (PAL) method. (d) Flow cytometry histograms and quantification of SLBR-N levels on MDA-MB231; SLBR-N levels quantified by measurement of mean fluorescent intensity in confocal microscopy images. (e-g) Flow cytometry analysis of MAL-II (e), Siglec-15 (f), and SNA (g) levels on MDA-MB231. (h) Flow cytometry analysis of PAL, SNA, and MAL-II on breast cancer cell lines with different bone-metastatic potential cultured on tissue culture plastic. In panel **b** and **d**, results are the mean ± s.d. (*n* = 3) and statistical differences were determined using two-tailed unpaired t-test.

Sialylated core-II O-glycans were also upregulated as revealed by increased binding of SLRB-N, which preferentially interacts with disialylated core-II O-glycans[66,69] (Fig. 5d). Higher SLBR-N reactivity correlated with higher Muc1 expression in the cells (Supplementary Fig. 4b-d). We also observed strong cell-surface reactivity with the MAL-II lectin, which preferentially recognizes the monosialyl-T and disialyl-T core-I O-glycan structures[70] (Fig. 5e). Siglec-15 has been reported to specifically bind the sialyl-Tn antigen mucins and regulate osteoclast differentiation from hematopoietic progenitors in the bone niche[71,72]. Therefore, we probed cells with recombinant Siglec-15, which showed a small increase in sialyl-Tn antigen levels in cells cultured on mineralized collagen (Fig. 5f). A larger increase was observed for α2,6-linked sialic acids probed by the Sambucus nigra lectin (SNA), which has binding preferences to α2,6 sialyl N-acetyllactosamine (LacNAc) on both N– and O-glycans[70] (Fig. 5g). Bone-metastatic BoM-1833 also exhibited a significantly higher expression of α2,6-linked sialic acids compared to the parental MD-MB231 cell line, suggesting that metastatic cells with a preference for bone metastasis formation may have unique sialic acid profiles. (Fig. 5h). We also evaluated the cell surface glycosylation levels in MCF7 cells and observed higher levels of a2,3-linked sialic acid and disialylated core-II O-glycans in the cells cultured on mineralized collagen substrates (Supplementary Fig. 3). Cell surface HA levels were elevated in both MDA-MB231 and MCF7 cells cultured on mineralized collagen, but the increased HA in the glycocalyx did not appear to significantly alter cell susceptibility to NK cell-mediated cytotoxicity, whereas mucin removal from the cancer cells with StcE enhanced their killing (Supplementary Fig. 3, 5). Overall, our findings implicate collagen mineralization in immune protection through the upregulation of mucin-type O-glycosylation and sialylation.

### 2.6. Inhibition of sialylation improves NK cell cytotoxicity of breast cancer cells interacting with mineralized collagen

We next tested whether the glycocalyx structure of mucin-expressing cells could be disrupted with a small molecule inhibitor of sialylation as a potential strategy to improve the efficacy of immunotherapies targeting bone metastasis (Fig. 6a). As an initial model, we measured the nanoscale thickness of the glycocalyx in a clonal MCF10A cell line that expresses Muc1-GFP under the control of a doxycycline inducible promoter. Prior studies have shown that the Muc1 expression levels in the clonal cell line, referred to as 1E7 cells, could be tuned through titration of the doxycycline concentration for induction[41]. With increased doxycycline concentrations, we observed marked increases in the glycocalyx thickness corresponding to graded increases in Muc1 and sialic acid surface levels (Fig. 6b,c). Targeting cell-surface sialylation using P-3F_AX_-Neu5Ac, a cell-permeable sialic acid inhibitor, disrupted mucin sialylation and dramatically reduced the glycocalyx thickness[73,74] (Fig. 6b,c). Compared to untreated control cells, the glycocalyx thickness of cells treated with P-3F_AX_-Neu5Ac was reduced by approximately 30 nm for cells expressing high levels of Muc1 (Fig. 6c). These results indicated that sialylation could be targeted to block the thickening of the glycocalyx resulting from the overexpression of cancer-associated mucins.

**Figure 6.**
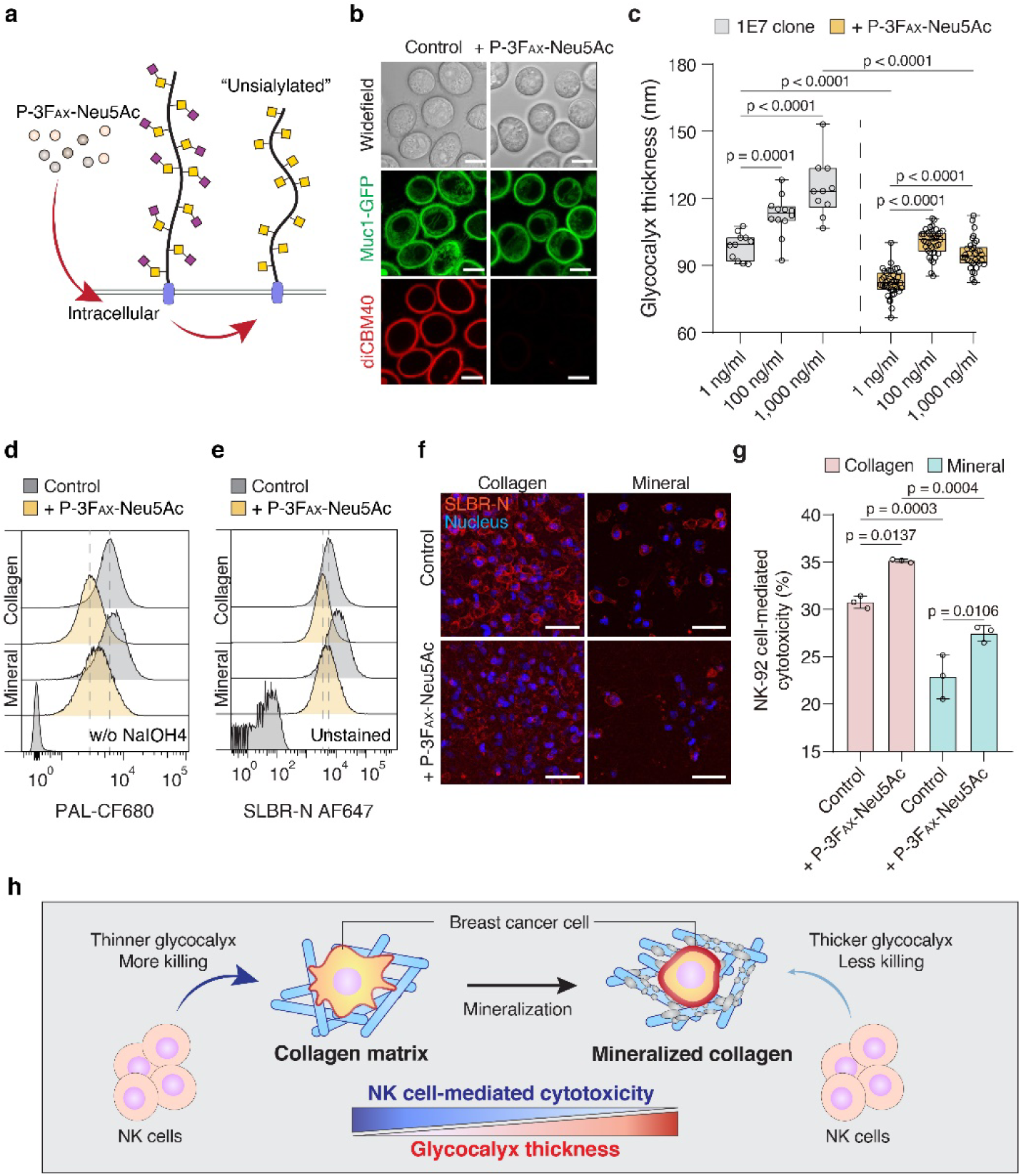
Small-molecule inhibition of cancer-cell sialylation enhances NK-92 cell-mediated cytotoxicity. (a) Schematic of sialic acid removal from the cell surface with the sialylation inhibitor – P-3F_AX_-Neu5Ac. (b) Representative confocal images of Muc1-GFP overexpressing MCF10A epithelial cells (1E7 clone) treated with DMSO or 100 µM sialylation inhibitor for 24 hours. Cell-surface α2,3 sialic acids were detected using Alexa Flour 647-labeled recombinant diCBM40. Scale bars, 10 µm. (c) Quantification of mean glycocalyx thickness in 1E7 cells whose expression of Muc1-GFP levels was controlled by varying the concentration of doxycycline before and after treatment with 100 µM sialylation inhibitor. Results are the mean ± s.d. of at least 10 cells. The bottom whisker represents the minimum, the top whisker represents the maximum and the square in the box represents the mean. (d,e) Representative flow cytometry histograms showing the effects of P-3F_AX_-Neu5Ac on MDA-MB231; sialylation probed using the PAL method (d) and SLBR-N (e). (f) Representative confocal images of MDA-MB231 cells cultured on collagen and mineralized collagen following treatment with 100 µM sialylation inhibitor for 24 hours. Scale bars, 50 µm. (g) NK-92 cell-mediated cytotoxicity against MDA-MB231 cells pretreated with 100 µM sialylation inhibitor for 24 hours. Results are the mean ± s.d. (*n* = 3). (h) Proposed relationship between bone matrix mineralization and glycocalyx-mediated immune evasion by breast cancer cells. In panel **c** and **g**, statistical differences were determined using one-way ANOVA with multiplicity-adjusted p values from Tukey’s multiple comparisons test.

Our results in the mucin-overexpressing cell model raised the possibility that sialylation could similarly be inhibited in cancer cells to circumvent the ability of mineralized collagen to support immune evasion of breast cancer cells through regulation of the glycocalyx structure. To test this possibility, MDA-MB231 cancer cells were cultured on collagen and mineralized collagen substrates and subsequently treated with P-3F_AX_-Neu5Ac. Within 24 hours of treatment with the inhibitor, the total levels of sialic acid on the cell surface were reduced to approximately 50% of their pretreatment levels (Fig. 6d). We confirmed that mucin sialylation also was specifically disrupted by the inhibitor in a similar timeframe. Sialylated core-2 O-glycans were probed with SLBR-N lectin and analyzed using flow cytometry and confocal microscopy (Fig. 6e,f). Mirroring results for total cell surface sialylation, SLBR-N binding was reduced by approximately 50% following inhibitor treatment of cells on collagen and mineralized collagen. These results align with prior reports similarly finding that sialylation inhibitors may not completely remove all sialic acid content within cells[74]. To confirm that the reduced sialylation abrogated the protective functioning of the glycocalyx, NK-92 cell-mediated cytotoxicity against the cancer cells was assayed (Fig. 6g). Cancer cells treated with P-3F_AX_-Neu5Ac exhibited increased susceptibility to killing by the NK-92 cells. Notably, the effect of inhibitor treatment was statistically significant even though the sialic acid content was only partially reduced over the treatment timeframe (Fig. 6g). Overall, our results indicate that small-molecule inhibitors that target cell-surface sialylation could serve as a promising strategy for improving current cell-mediated immunotherapies in the context of bone-metastatic breast cancer (Fig. 6h).

## 3. Discussion

The rapid advance of anticancer immunotherapy into clinical practice underlies the urgent need to identify the molecular mechanisms of immune evasion within the context of specific microenvironments. Identifying these connections is particularly important for patients with advanced breast cancer as their tumor cells often disseminate to bone, where they can adopt latent and immune-protected phenotypes that can later develop into overt metastasis that are challenging to treat. Our findings suggest that collagen mineralization may play a pivotal role in protecting DTCs within skeletal niches by favoring the synthesis of a thicker cancer-cell glycocalyx that can support evasion of NK cell-mediated cytotoxicity. Notably, our work reveals mucins and sialoglycans as key glycocalyx structural elements that may protect against the onslaught of effector immune cells. As such, therapeutic targeting of sialylation or cell-surface mucins could represent a possible strategy to improve the efficacy of immunotherapies in the context of bone-metastatic breast cancer, but further experiments are necessary to validate our findings *in vivo*.

By design, our studies with engineered collagen substrates were intended to isolate the effect of bone matrix mineralization on the glycocalyx and evasion of NK-cell mediated cytotoxicity. We were interested in these connections because tumor cells interact with bone matrix following dissemination to osteogenic niches[6] and because bone mineral density decreases with various conditions associated with increased risk for bone metastasis including age, vitamin D deficiency, and chemotherapy[75–77]. The ability of the bone matrix to regulate the glycosylation of disseminated tumor cells could have broad consequences within the full complexity of the bone microenvironment. Of the many possibilities, we observed elevated levels of α2,6 sialic acids in the bone-homing BoM-1833 cancer line and when breast cancer cells were cultured on mineralized collagen. Elevated levels of α2,6 sialic acid on tumors cells is associated with poorer effector T cell responses, reduced tumor infiltration of NK cells, and lower NK cell activity[78]. Sialic acid also plays an important role in the differentiation of osteoclasts and the interaction between osteoclasts and other cells. In particular, α2,6 sialic acids are strongly implicated in the regulation of osteoclast differentiation from hematopoietic precursors[72,79,80]. During late-stage bone metastasis, osteoclastogenesis and osteolytic bone resorption set off a positive feedback loop, referred to as the “vicious cycle,” in which bone degradation releases pro-tumor factors from the bone matrix to promote further tumor cell growth and stimulation of bone remodeling[81]. Hence, it is possible that tumor cells interacting with mineralized bone matrix use glycocalyx changes as a mechanism to induce osteoclastogenesis, which stimulates bone matrix changes that activate tumor growth over time. Interestingly, we have found previously that interactions with mineral induce a stem-like, more latent phenotype in tumor cells whose gene expression correlates with improved patient prognosis but was reversible when bone mineral content decreased[7]. As immune evasion is a hallmark of latent cancer stem-like cells[16] and because matrix properties can permanently change cell phenotype[82], it is conceivable that tumor cells lastingly reprogram their expression of glycan genes when interacting with mineralized bone matrix to evade immune attack even under conditions of reduced bone mineral density during later stages of the disease. In lieu of these connections, our work supports further investigation of how altered DTC sialylation in mineralized osteogenic niches may reshape the broader cellular microenvironment and how these collective changes regulate bone metastasis and long-term patient prognosis.

The correlation that we observed between reduced NK-cell mediated cytotoxicity and the nanoscale thickness of the cancer cell glycocalyx across varying matrix conditions is striking and adds to the growing body of evidence implicating the biophysical properties of the glycocalyx as major determinants of immune evasion[41,83–86]. In addition to the glycocalyx thickness, the compressibility of the glycocalyx layer has long been predicted to resist the specific interactions of cells[35]. Experimental tools that can measure the elasticity and other material properties of the glycocalyx must still be developed. However, theoretical studies have shown that the deformation of the glycocalyx under compressive stress could vary considerably and in highly nonlinear ways depending on whether the mucin constituents have a flexible versus more rigid, rod-like structure[87]. These molecular properties depend on the density and profile of O-glycans along the mucin polypeptide backbone, raising the possibility that changes in glycosylation linked to specific microenvironments, like bone, may regulate the physical properties of individual mucins or other glycoproteins to modulate the deformability of the glycocalyx layer.

With the explosion of interest in tumor glycobiology over the past decade, multiple strategies for targeting the glycocalyx have been developed, some of which have advanced to human clinical testing. Here, we show that small molecule drugs that inhibit sialylation may be beneficial in mitigating immune evasion within osteogenic niches. Although not tested in this work, small-molecule inhibitors of mucin-type O-glycosylation also have demonstrated *in vivo* safety and functionality, raising the possibility that they can be used alone or in combination with sialylation inhibitors to therapeutically target the glycocalyx structure[88]. To improve the specificity of targeting, antibody-enzyme conjugates have been developed that direct mucinases or sialidases toward a target antigen on the cancer cell surface to disrupt the glycocalyx structure for improved immunotherapy[89–91]. Furthermore, our recent work has shown that glycocalyx-modifying enzymes, like mucinases can be tethered directly to the NK cell surface as another potential strategy to improve adoptive cell therapies[41]. The findings here support continued development and evaluation of such strategies to improve the performance of immunotherapy within challenging microenvironments, like bone.

While our results suggest clear functional links between tumor cell phenotype, collagen mineralization, glycocalyx formation, and immunoregulation, future experiments will need to elucidate in more detail how these parameters regulate tumor cell latency and metastasis at skeletal sites. For example, other microenvironmental factors such as culture dimensionality (2D vs 3D) and co-culture with different bone niche cells can also affect the fate of DTCs[92,93], but have not been considered in the described experiments. Expanding our culture platforms to recapitulate some of these parameters will enhance their relevance and may provide additional insights into how collagen mineralization regulates tumor cell latency and immunoregulation under more complex microenvironmental conditions. Additionally, it will be important to study T-cell responses in these varied contexts and perform *in vivo* studies in immunocompetent animals to investigate how bone matrix density regulates tumor-immune cell interactions and relate these findings to differences in glycocalyx biosynthesis.

## 4. Conclusion

In this study, we revealed a significant and inverse relationship between the bone metastatic potential of breast cancer cells and their physical glycocalyx thickness in the context of NK cell-mediated cytotoxicity. We demonstrate that collagen mineralization plays a pivotal role in modulating the immunoregulatory processes in breast cancer cells, leading to increased glycocalyx thickness and enhanced evasion of NK attacks. Notably, mucins and sialoglycans emerge as key protective factors for cancer cells against the onslaught of effector immune cells. Furthermore, our findings highlight the regulatory role of collagen mineralization in mucin sialoglycans. Importantly, the inhibition of sialylation emerges as a promising strategy to enhance NK cell cytotoxicity against breast cancer cells in bone matrices. These results offer valuable insights into the development of therapeutic approaches to cancer immunotherapies in the context of bone-metastatic breast cancer by targeting the cancer glycocalyx and its interactions with the immune system.

## 5. Materials and methods

### 5.1. Cell culture

MDA-MB231 (ATCC), MCF7 (ATCC) and bone metastatic BoM-1833 breast cancer cell lines (kindly provided by Joan Massague) were cultured in Dulbecco’s Modified Eagle Medium (DMEM, Thermo Fisher Scientific) supplemented with 10% fetal bovine serum (FBS, Atlanta Biologicals) and 1× penicillin/streptomycin (Thermo Fisher Scientific) at 37°C in 5% CO_2_.

NK-92 cells (ATCC; CRL-2407) were cultured in Minimum Essential Medium α (_-MEM) without ribonucleosides media (Thermo Fisher Scientific) supplemented with 12.5% fetal bovine serum, 12.5% horse serum, 0.2 mM Myo-inositol (Sigma Aldrich), 0.1 mM 2-mercaptoethanol (Thermo Fisher Scientific), 0.02 mM folic acid (Millipore Sigma), 100 U/mL recombinant human IL-2 (Pepro Tech), and 1× penicillin/streptomycin at 37°C in 5% CO_2._

A clonal MCF10A cell line expressing Muc1-GFP was generated previously and referred to here as 1E7 cells[41]. The cells were cultured in DMEM/F12 media (Thermo Fisher Scientific) supplemented with 5% horse serum (Thermo Fisher Scientific), 20 ng/mL EGF (Pepro Tech), 10 mg/mL insulin (Sigma), 500 ng/mL hydrocortisone (Sigma), 100 ng/mL cholera toxin (Sigma) and 1x penicillin/streptomycin (Thermo Fisher Scientific) at 37°C in 5% CO_2_.

### 5.2. Isolation and culture of primary human NK cells

Human peripheral blood mononuclear cells (PBMCs) were isolated from human buffy coats from the New York Blood Center by density gradient centrifugation using Ficoll-Paque PLUS (GE Healthcare). Primary NK cells were extracted from PBMCs by negative immunomagnetic isolation using a NK cell isolation kit (Miltenyi Biotec) according to the manufacturer’s protocol. Briefly, cells were incubated with NK Cell Biotin-Antibody Cocktail and NK Cell MicroBead Cocktail to label non-NK cells, followed by loading onto a LS Separation column (Miltenyi Biotec). The cell suspension was then placed in the magnetic field of a MACS Separator to separate labeled from non-labeled cells. The isolated human NK cells were subsequently cultured in Roswell Park Memorial Institute (RPMI) 1640 media supplemented with 10% fetal bovine serum, 1x penicillin/streptomycin and 100 U/mL IL-2 at 37°C in 5% CO_2_. The purity of the NK cell population was validated by flow cytometry using PE-Vio 770 anti-human CD56 (1:50; REA196 clone; Miltenyi Biotec) and FITC anti-human CD3 (1:50; REA613 clone; Miltenyi Biotec) antibodies.

### 5.3. Fabrication and characterization of mineralized collagen

Fibrillar collagen substrates were mineralized using the polymer-induced liquid-precursor (PILP) process in polydimethylsiloxane (PDMS, Dow) microwells (diameter: 4 mm, height: 250 µm) or on glass cover slips as previously described[7,15,94]. PDMS microwells were fabricated using conventional soft-lithography methods and used for all imaging-related applications while the glass coverslip platform was used for all applications that require large cell numbers, including flow cytometry, SAIM, and cytotoxicity assay. To prepare cover slip substrates, 18 mm-diameter disks were punched out of cured approximately 1.5 mm thick PDMS sheets and adhered to 18 mm-diameter glass coverslips (VWR) using optical adhesive (Norland Products) under UV exposure. Both PDMS microwells and glass surfaces were treated with 1% (w/v) polyethyleneimine (PEI, Sigma-Aldrich) in distilled water and 0.5% glutaraldehyde (v/v) in distilled water (Sigma-Aldrich) to allow covalent binding of cast collagen type I (1.5 mg/mL, adjusting pH to 7.4 using 0.1N NaOH, Corning). After 2 hours of gelation at room temperature under humidified conditions, the formed collagen hydrogels were mineralized using a solution of 62.5 µg/mL of poly aspartic acid (PAA, MW = 27 kDa, Alamanda Polymers), 1.67 mM CaCl_2_ (Thermo Fisher Scientific) and 1 mM (NH_4_)_2_HPO_4_ (Sigma-Aldrich) in 0.85× phosphate buffered saline (PBS). This incubation was done in an upside-down configuration at 37°C for 2 days ensuring mineralization initiated within rather than on top of collagen fibrils. Collagen control substrates were prepared similarly but incubated in PBS rather than mineralizing solution. Mineralization of collagen was confirmed by scanning electron microscopy (SEM) and backscattered electron (BSE) imaging (Mira3 LM, Tescan). To this end, samples were fixed with 2.5% glutaraldehyde (0.05 M cacodylate, pH 7.4) and dehydrated using a series of ethanol solutions followed by incubation in hexamethyldisilazane (Electron Microscopy Sciences) and coated with carbon using a sputter coater (Desk V, Denton Vacuum). BSE images were converted to pseudocolour images to show differences in mineral content using MATLAB (MathWorks).

### 5.4. Characterization of cell adhesion and growth

Substrate-induced changes of cell morphology were analyzed 5 hours after seeding. For SEM imaging, samples were fixed and then prepared and imaged as described above. For cell morphology analysis, cells were fixed with 4% formaldehyde, and stained with DAPI and Alexa Fluor 488 phalloidin (Thermo Fisher Scientific). Images were captured as z-stacks with a 5 µm-step size using a confocal microscope (710 LSM, Zeiss) and a 40× water immersion objective (C-Apochromat, 1.2 NA). The collagen structure was imaged with confocal reflectance microscopy using a laser at 488 nm on the Zeiss 710 LSM. ImageJ was used to analyze the projected area of all adherent cells in 6 randomly selected images per sample (*n* = 3). Substrate-induced changes of cell proliferation were analyzed after 5 days of culture using the Click-iT EdU Alexa Fluor 647 Imaging Kit (Thermo Fisher Scientific) according to the manufacturer’s protocol. Briefly, cells were incubated with EdU for 4 hours before fixation with 4% formaldehyde. Fixed cells were permeabilized with 0.5% Triton X-100 in PBS, labeled with EdU Alex Fluor 647 for 30 minutes at room temperature, and counterstained with Hoechst dye. EdU positive cells were quantified using ImageJ in 5 randomly selected images per sample (*n* = 3).

### 5.5. RNA-seq data analysis

To assess transcriptomic changes of glycocalyx-associated genes induced by mineralized collagen, previously published RNA-sequencing results comparing MDA-MB231 cells cultured on mineralized or unmineralized collagen were retrieved from the Gene Expression Omnibus repository (GSE229094). Gene counts were analyzed using DESeq2[95] with differential genes defined as genes with an adjusted P value ≤ 0.05 and | log_2_(Fold-change) | ≥ 0.3. For heatmaps normalized gene counts calculated by DESeq2 were extracted, which account for differences in RNA library size between samples. Genes were categorized as glycosyltransferases, glycoside hydrolases, or other human glycan genes using previously annotated gene sets[96]. Within heatmaps hierarchical clustering of both individual genes and samples used the Euclidean distance metric.

### 5.6. Cell harvest and treatment with glycocalyx editing reagents

Tumor cells were detached from the different substrates after 7 days of culture. Cells cultured on tissue culture polystyrene were collected using 0.25% trypsin-EDTA. Cells cultured on collagen and mineralized collagen were collected using collagenase type I (1mg/mL in PBS, Worthington Biochemical) and passed through a 40 µm cell strainer. MCF10A-derived 1E7 cells cultured on tissue culture polystyrene were induced with 1 μg/mL doxycycline (204734, Santa Cruz Biotechnology) for 24 hours before being collected with 0.05% trypsin-EDTA.

For treatment with glycocalyx-editing enzymes, detached cells were incubated with 100 nM StcE mucinase or 100 nM *Salmonella typhimurium* sialidase or 2.5 U/mL *Streptomyces hyalurolyticus* hyaluronidase (H1136, Millipore Sigma) or 5,000 U/mL PNGase F (P0704S, New England BioLabs) in growth media for 1 hour at 37°C. Subsequently, cells were rinsed thoroughly twice with growth media and used further. For treatment with the sialylation inhibitor P-3F_AX_-Neu5Ac, tumor cells were cultured on collagen and mineralized collagen for 6 days before incubation with 100 μM P-3F_AX_-Neu5Ac in growth media for another 24 hours at 37°C in 5% CO_2_. Subsequently, cells were collected using collagenase type I (1mg/mL in PBS, Worthington Biochemical) and passed through a 40 µm cell strainer before use in subsequent assays.

### 5.7. Scanning angle interference microscopy (SAIM)

Silicon wafers with a ∼2,000 nm thermal oxide layer (Addison Engineering) were diced into 7 × 7 mm chips, and the oxide layer thickness of each chip was measured with a FilMetrics F50-EXR. Silicon chips then were functionalized using 4% (v/v) (3-mercaptopropyl)trimethoxysilane in absolute ethanol for 30 minutes at room temperature, followed by incubation with 4 mM 4-maleimidobutyric acid N-hydroxysuccinimide ester in absolute ethanol and 50 µg/ml human Alexa Fluor 647 conjugated plasma fibronectin as previously reported[41,45]. Cells were seeded onto the fibronectin-coated chip at 0.5-1_×_10^5^_cells/cm^2^ in full culture medium. After 4-6 hours, adhered cells were rinsed with serum-free, phenol-red free DMEM and incubated with MemGlow dyes (MemGlow 560, MG02-2 and MemGlow 590, MG03-10; Cytoskeleton) in serum-free, phenol red-free DMEM for 10 minutes at 37°C. Cell-seeded chips were then washed with serum-free, phenol red-free DMEM again, and inverted onto a 35 mm glass-bottom imaging dish and imaged at 37°C. As previously reported[41], SAIM was conducted on a custom circle-scanning microscope which allowed imaging at varying incidence angles, ranging from 5 to 43.75 degrees, and a total of 32 images was acquired per cell. The intensities of raw image sequences were fit pixelwise by nonlinear least-squares to an optical model:

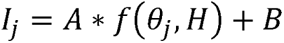

where *I_j_* is raw image intensity at each incidence angle *θ_j_*, H is the glycocalyx thickness, and A and B are additional fit parameters. The optical system maintained the s-polarization of circle-scanned excitation laser by the vortex half-wave plate. The probability of excitation is given by:

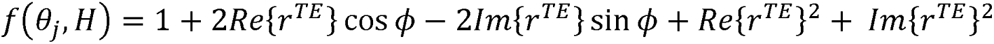

Where □(*H*) is the phase shift, λ is s-polarized monochromatic excitation of wavelength, and *r^TE^* is the reflection coefficient for the transverse electric wave and these are given by:

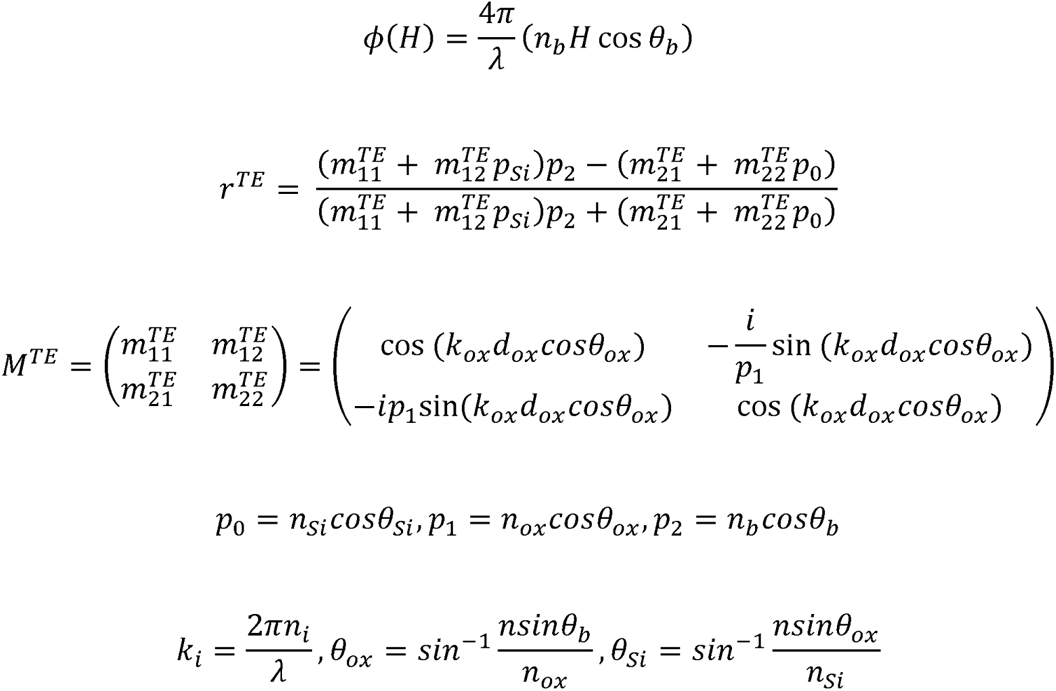

where *k_i_* is the wavenumber in material i; *n_Si_*, *n_ox_* and *n_b_* are the refractive index of the silicon, silicon oxide and sample, respectively; *θ_Si_*, *θ_ox_* and *θ_b_* are the angles of incidence in the silicon, silicon oxide and sample, respectively; and *d_ox_* is the thickness of the silicon oxide layer. The angles of incidence in silicon oxide and silicon were calculated according to Snell’s Law. The average glycocalyx thickness was quantified in 200 × 200 pixel subregions of each cell by subtracting the height of their MemGlow signal from the height of the corresponding fluorescently labeled fibronectin on the silicon substrate.

### 5.8. Primary NK & NK-92 cell-mediated cytotoxicity assays

Detached cells were fluorescently labeled for 15 minutes with 10 μM CellTracker Green CMFDA Dye (Invitrogen) in growth media, followed by washing thoroughly twice with growth media. 4×10^4^ labeled tumor cells were mixed with varying ratios of NK-92 cells or primary NK cells in 200 µL of growth media of the tumor cell line in the absence of IL-2 and co-cultured in an ultra-low attachment U-bottom 96-well plate (Corning) for 4 hours at 37°C in 5% CO_2_. Following the 4-hour co-culture, the mixed cells were pelleted by centrifugation at 500 g for 5 minutes, resuspended and incubated in propidium iodide solution (PI; 20 µg/mL, Sigma) for 10 minutes. NK cell-mediated cytotoxicity was then measured by flow cytometry as previously described[41,42,97]. At least 1 ×10^4^ tumor cells were analyzed after electronic gating on CellTracker Green. To calculate the percent cytotoxicity, the following formula was used: 100 × (experimental % dead − spontaneous % dead)/(100 − spontaneous % dead), where experimental % dead was the percentage of PI positive tumor cells in co-cultures and spontaneous % dead was the percentage of PI positive control tumor cells cultured in the absence of effector cells.

### 5.9. Preparation of recombinant StcE mucinase and *Salmonella typhimurium* sialidase

The cDNA for StcE-Δ35 and sialidase was synthesized by custom gene synthesis (Twist Bioscience) and inserted into the pET28b expression vector[37]. The recombinant enzymes were produced by expressing them in chemically competent NiCo21 (DE3) E. coli (NEB). Following transformation and overnight growth on Luria Broth (LB) agar plates, the cells were cultured in LB medium at 37°C until an OD600 of 0.6-0.8 was reached. At this point, the cultures were induced with 0.5 mM isopropyl β-D-1-thiogalactopyranoside (IPTG) and grown overnight at 24°C. Cells were then harvested by centrifugation at 3,000 g for 20 minutes, resuspended in lysis buffer (20 mM HEPES, 500 mM NaCl and 10 mM imidazole, pH 7.5) with cOmplete protease inhibitor Cocktail (Roche), and lysed by a sonicator (Q125, Qsonica). Recombinant enzymes were purified by immobilized metal affinity chromatography (IMAC) on a GE ÄKTA Avant FPLC system. The lysate was applied to a HisTrap HP column (Cytiva), followed by a wash step with 20 column volumes of wash buffer (20 mM HEPES, 500 mM NaCl and 20 mM imidazole, pH 7.5). Elution was performed using a linear gradient of 20 mM to 250 mM imidazole in buffer (20 mM HEPES and 500 mM NaCl, pH 7.5). The eluted fractions containing target protein were collected and further refined by a HiPrep 26/60 Sephacryl S-200 HR (Cytiva) column equilibrated with storage buffer (20 mM HEPES and 150 mM NaCl, pH 7.5). The final protein was concentrated by using Amicon Ultra 30 kDa MWCO filters (Millipore Sigma).

### 5.10. Preparation of recombinant diCBM40 and conjugation[99]

The cDNA for diCBM40 from *Clostridium perfringens* was synthesized N-terminal 6x-His tag by custom gene synthesis (General Biosystems) and inserted in pET21 expression vector. diCBM40 was produced by expressing it in chemically competent NiCo21 (DE3) E. coli (NEB) as described above for StcE and sialidase. The harvested cells were then lysed using B-PER (Thermo Fisher Scientific), and the lysates were cleared by centrifugation at 10,000 g for 20 minutes at 4°C. The diCBM40 was purified by IMAC using spin columns according to standard protocols. In brief, the clarified lysate was diluted into 1x Ni-NTA binding buffer and bound to equilibrated Ni-NTA resin (Qiagen, 30210) for 20 minutes at 4°C with end-over-end mixing. Subsequently, the resin was packed into a spin column, washed thoroughly, and then incubated with the Ni-NTA elution buffer for 20 minutes at 4°C, mixing end-over-end. Eluted protein was exchanged into storage buffer (PBS, pH 7.4) using Zeba 7K MWCO desalting columns. Eluted proteins were then sterile filtered and snap-frozen for long-term storage at –80°C. For conjugation with fluorophore, purified diCBM40 was labeled with Alexa Fluor 647 NHS Ester (Thermo Fisher Scientific) per the manufacturer’s protocol.

### 5.11. Preparation of recombinant SLBR-N and conjugation

The cDNA for SLBR-N in a pGEX expression vector was obtained as a gift of Barbara Bensing[100]. SLBR-N was recombinantly produced in chemically competent NiCo21 (DE3) E. coli (NEB) as described above for StcE and sialidase with the exception that expression was induced with IPTG at an OD600 of 0.9 The harvested cells were then lysed using sonication, and the lysates were cleared by centrifugation at 10,000 g for 20 minutes at 4°C. The GST-tagged SLBR-N was purified using Glutathione Sepharose 4B resin (Cytiva) with gravity flow columns according to the manufacturer’s instructions. Eluted protein was exchanged into storage buffer (PBS, pH 7.4) using Zeba 7K MWCO desalting columns. For conjugation with fluorophore, purified SLBR-N was labeled with Alexa Fluor 647 NHS Ester (Thermo Fisher Scientific) per the manufacturer’s protocol.

### 5.12. Lectin staining

Detached target breast cancer cells were incubated with different fluorescently labelled or biotin-labelled lectins (PNA-CF650R; HABP-biotin; diCBM40-Alexa Fluor 647; SLBR-N Alexa Fluor 647; SNA-Cy5, MAL-II-biotin) at 4°C for 1 hour. Lectins were diluted 1:200 in 0.5% BSA PBS and incubated with cells at 4°C for 1 hour. Secondary labelling was with NeutrAvidin-DyLight 650 (84607; Thermo Fisher Scientific) or Streptavidin-Alexa Fluor 488 (S11223; Thermo Fisher Scientific), diluted 1:200 in 0.5% BSA PBS and incubated with cells at 4°C for 1 hour. The Attune NxT flow cytometry (Thermo Fisher Scientific) was used for analysis.

For detecting diCBM40 on 1E7 cells, cells were plated in 35 mm glass bottom dishes (P35G-1.5-14-C; Matteck), grown for 24 hours, and then induced with 1,000 ng/ml of doxycycline for 24 hours. Cells were then incubated with diCBM40 lectins at 4°C for 1 hour. After washing with 0.5% BSA PBS, cells were imaged on a LSM 800 confocal microscopy using 63× (NA: 1.2 Water) objectives (Zeiss). For detecting SLBR-N binding on MDA-MB231, the cells were plated in 35 mm glass bottom dishes (P35G-1.5-14-C; Matteck), grown for 24 hours. Cells were then incubated with SLBR-N conjugated Alexa Fluor 647 at 4°C for 1 hour. After washing with 0.5% BSA PBS, cells were imaged on a LSM 800 confocal microscopy using 20× (NA: 0.8 Air) objectives (Zeiss). For detecting other lectin bindings on MDA-MB231, the cells plated on collagen or collagen mineralization and grown for 7 days. Images were captured using a confocal microscope (710 LSM, Zeiss) and a 40× water immersion objective (C-Apochromat, 1.2 NA). ImageJ was used for analysis.

### 5.13. Periodate labelling of cell surface sialic acid[68]

Detached cells were incubated with 1 mM sodium periodate in PBS at 4°C for 30 minutes. The periodate was quenched by 1 mM glycerol in cold PBS, followed by washing with cold PBS. The cells were then incubated with 25 µM Alexa Fluor 568-hydroxylamine (Fluoroprobes) or CF680R-Aminooxy (Biotium) in the presence of 10 mM aniline (Sigma) in 5% FBS PBS pH 6.7 at 4°C for 1 hour. Subsequently, the cells were rinsed twice with cold 0.5% BSA PBS. Flow cytometry was performed on an Attune NxT flow cytometry (Thermo Fisher Scientific).

### 5.14. Statistical analysis

Unless otherwise indicated, results are presented as the mean and standard deviation (s.d.) of at least 3 replicates per condition using GraphPad Prism 9. Statistical differences were determined using two-tailed unpaired t-test for two group comparisons, one-way ANOVA with multiplicity-adjusted p values from Tukey’s multiple comparisons test and Newman-Keuls’ multiple comparisons test. Two-tailed test with p < 0.05 were considered significant in all cases.

## Supporting information

Supproting Information

## 6. Acknowledgment

We thank J. Massague (Memorial Sloan Kettering Cancer Center) for providing bone tropic BoM-1833. This investigation was supported by National Science Foundation (NSF) grant 1752226 (M.J.P.), the Breast Cancer Coalition of Rochester predoctoral fellowship (S.P.), and National Institutes of Health (NIH) grants: R01 CA259195 (C.F.), U54 CA210184 (C.F. and M.J.P.), R01 CA276398 (M.J.P.), DP2 GM229133 (M.J.P.), R01 GM138692 (M.J.P).

Work was performed at the Center for Materials Research (CCMR), which is supported through the NSF MRSEC program (DMR-1719875); the Cornell Nanoscale Facility (NSF NNCI-2025233), a member of the NSF-supported National Nanotechnology Coordinated Infrastructure (NNCI-2025233); Biotechnology Resource Center (RRID:SCR_021740), and Imaging Facility (RRID:SCR_021741) with NYSTEM (C029155), NIH (S10OD018516), NIH (1S10RR025502) funding for the Zeiss LSM880 and LSM 710 confocal microscope. SAIM instrument development was supported by the Kavli Institute at Cornell for Nanoscale Science.

## 7. Author contributions

S.P., S.C., M.J.P., and C.F. designed the project. S.P., and S.C., conducted and analyzed all cytotoxicity assay and flow cytometry. S.C. performed mineralized matrix fabrication and analysis. S.P. conducted the SAIM measurements and analysis. A.A.S. analyzed RNA-seq data. S.P. S.C., M.J.P., and C.F. wrote the manuscript with feedback from all authors.

## 8. Author Information

Claudia Fischbach

Matthew J. Paszek

Address: 120 Olin Hall, 113 Ho Plaza, Cornell University, Ithaca NY 14853, USA

