## Supplementary material for "Collagen mineralization decreases NK cell-mediated cytotoxicity of breast cancer cells via increased glycocalyx thickness": Supproting Information

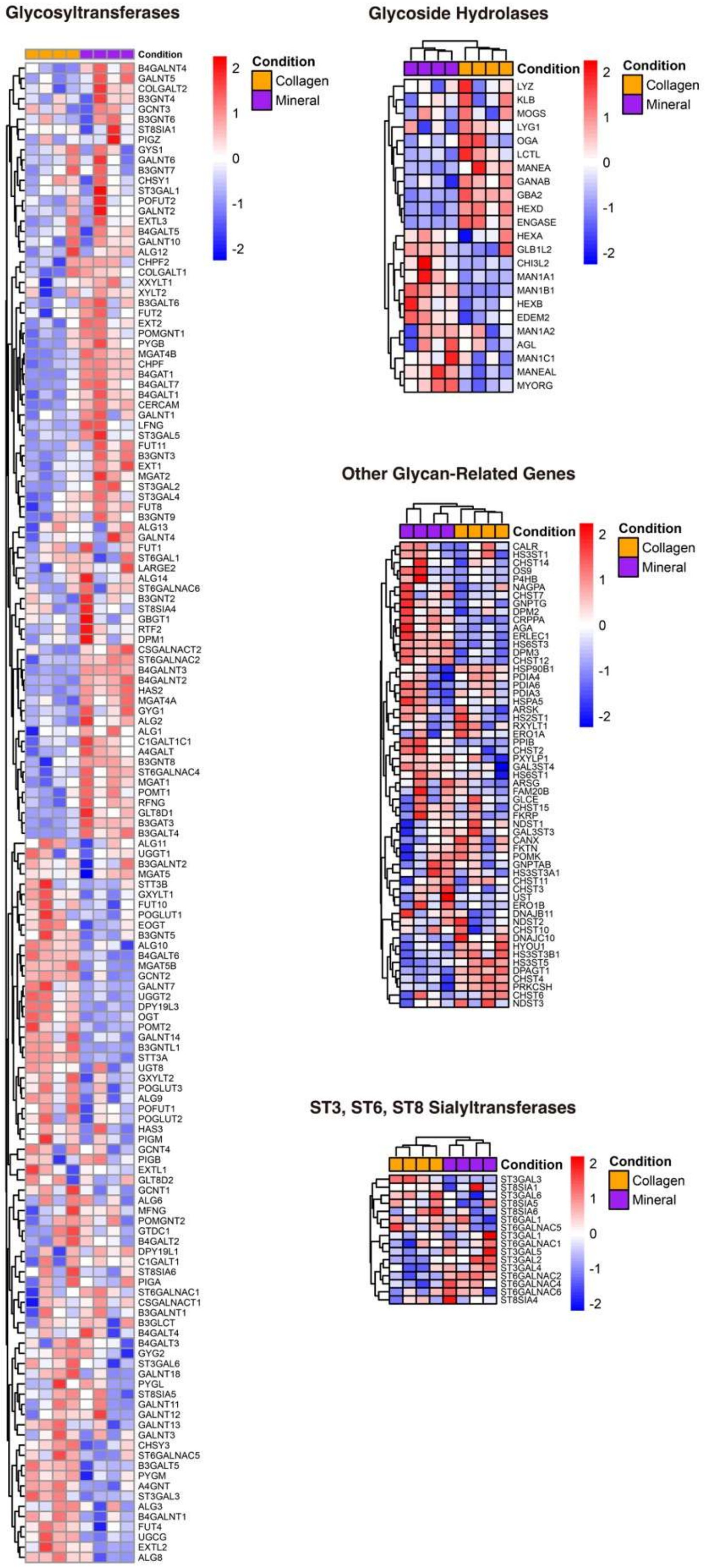
Glycosyltransferases
Condition
Collagen
Mineral
2
1
0
-1
-2
B4GALNT4
GALNT5
COLGALT2
B3GNT4
GCNT3
B3GNT6
ST8SIA1
PIGZ
GYS1
GALNT6
B3GNT7
CHSY1
ST3GAL1
POFUT2
GALNT2
EXTL3
B4GALT5
GALNT10
ALG12
CHPF2
COLGALT1
XXYLT1
XYLT2
B3GALT6
FUT2
EXT2
POMGNT1
PYGB
MGAT4B
CHPF
B4GAT1
B4GALT7
B4GALT1
CERCAM
GALNT1
LFNG
ST3GAL5
FUT11
B3GNT3
EXT1
MGAT2
ST3GAL2
ST3GAL4
FUT8
B3GNT9
ALG13
GALNT4
FUT1
ST6GAL1
LARGE2
ALG14
ST6GALNAC6
B3GNT2
ST8SIA4
GBGT1
RTF2
DPM1
CSGALNACT2
ST6GALNAC2
B4GALNT3
B4GALNT2
HAS2
MGAT4A
GYG1
ALG2
ALG1
C1GALT1C1
A4GALT
B3GNT8
ST6GALNAC4
MGAT1
POMT1
RFNG
GLT8D1
B3GAT3
B3GALT4
ALG11
UGGT1
B3GALNT2
MGAT5
STT3B
GXYLT1
FUT10
POGLUT1
EOGT
B3GNT5
ALG10
B4GALT6
MGAT5B
GCNT2
GALNT7
UGGT2
DPY19L3
OGT
POMT2
GALNT14
B3GNTL1
STT3A
UGT8
GXYLT2
POGLUT3
ALG9
POFUT1
POGLUT2
HAS3
PIGM
GCNT4
PIGB
EXTL1
GLT8D2
GCNT1
ALG6
MFNG
POMGNT2
GTDC1
B4GALT2
DPY19L1
C1GALT1
ST8SIA6
PIGA
ST6GALNAC1
CSGALNACT1
B3GALNT1
B3GLCT
B4GALT4
B4GALT3
GYG2
ST3GAL6
GALNT18
PYGL
ST8SIA5
GALNT11
GALNT12
GALNT13
GALNT3
CHSY3
ST6GALNAC5
B3GALT5
PYGM
A4GNT
ST3GAL3
ALG3
B4GALNT1
FUT4
UGCG
EXTL2
ALG8
Glycoside Hydrolases
Condition
Collagen
Mineral
2
1
0
-1
-2
LYZ
KLB
MOGS
LYG1
OGA
LCTL
MANEA
GANAB
GBA2
HEXD
ENGASE
HEXA
GLB1L2
CHI3L2
MAN1A1
MAN1B1
HEXB
EDEM2
MAN1A2
AGL
MAN1C1
MANEAL
MYORG
Other Glycan-Related Genes
Condition
Collagen
Mineral
2
1
0
-1
-2
CALR
HS3ST1
CHST14
OS9
P4HB
NAGPA
CHST7
GNPTG
DPM2
CRPPA
AGA
ERLEC1
HS6ST3
DPM3
CHST12
HSP90B1
PDIA4
PDIA6
PDIA3
HSPA5
ARSK
HS2ST1
RXYLT1
ERO1A
PPIB
CHST2
PXYLP1
GAL3ST4
HS6ST1
ARSG
FAM20B
GLCE
CHST15
FKRP
NDST1
GAL3ST3
CANX
FKTN
POMK
GNPTAB
HS3ST3A1
CHST11
CHST3
UST
ERO1B
DNAJB11
NDST2
CHST10
DNAJC10
HYOU1
HS3ST3B1
HS3ST5
DPAGT1
CHST4
PRKCSH
CHST6
NDST3
ST3, ST6, ST8 Sialyltransferases
Condition
Collagen
Mineral
2
1
0
-1
-2
ST3GAL3
ST8SIA1
ST3GAL6
ST8SIA5
ST8SIA6
ST6GAL1
ST6GALNAC5
ST3GAL1
ST6GALNAC1
ST3GAL5
ST3GAL2
ST3GAL4
ST6GALNAC2
ST6GALNAC4
ST6GALNAC6
ST8SIA4

**Supplementary Figure 1.** Heatmap of glycogenes (categorized as glycosyltransferases, glycoside hydrolases, sialyltransferases, and other glycan-related genes) differentially expressed by MDA-MB231 cells on collagen versus mineralized collagen. GEO accession number is GSE229094.

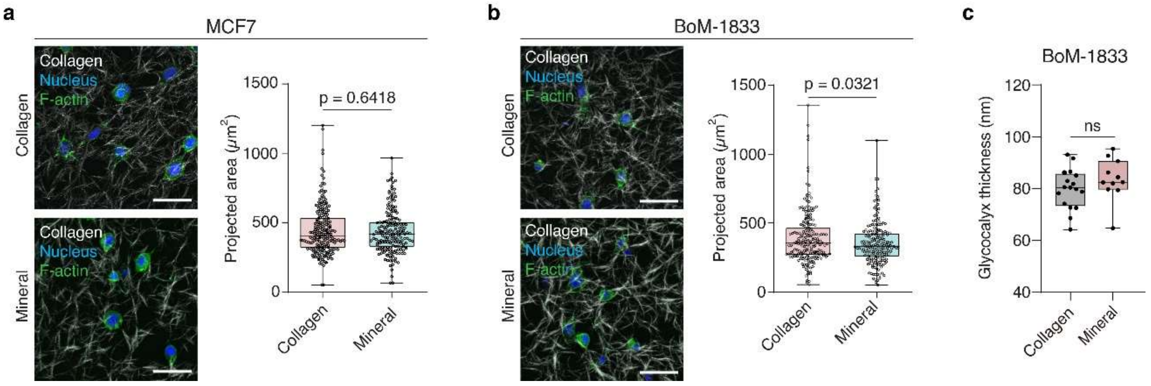


**Supplementary Figure 2. Effects of collagen mineralization on cell morphology of MCF7 and BoM-1833 and glycocalyx thickness of BoM-1833.** (a,b) Representative confocal images and corresponding quantification of cell morphology of MCF7 (a) and BoM-1833 (b) by image analysis. Scale bars, 50 µm. Boxes and whiskers show the first and third quartiles (boxes), median, and range of the data. Each condition includes a minimum of 202 cells from a representative experiment. In panel **a** and **b**, statistical analysis by two-tailed unpaired t-test with Welch's correction. (c) Quantification of glycocalyx thickness in BoM-1833 cells precultured on collagen and mineralized collagen. Boxes and whiskers show the first and third quartiles (boxes), median, and range of the data. Each condition includes a minimum of 11 cells from a representative experiment. Statistical differences were determined using two-tailed unpaired t-test.

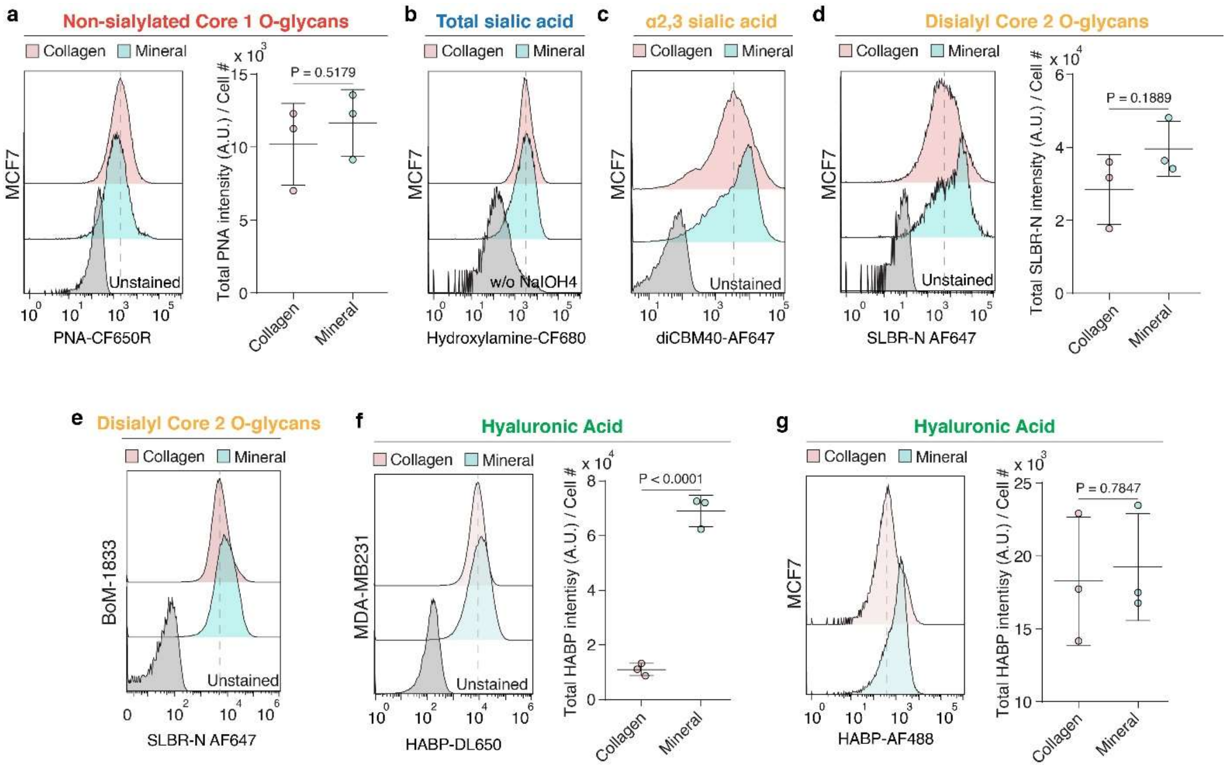


**Supplementary Figure 3. Collagen mineralization enhances the expression of sialoglycans and hyaluronic acid in MDA-MB231 and MCF7 cells.** (a) Flow cytometry histograms and quantification of PNA levels on MCF7 cells precultured on collagen or mineralized collagen (mineral); PNA levels quantified by measurement of mean fluorescent intensity in confocal microscopy images. (b) Flow cytometry analysis of total cell-surface sialic acids on MCF7 cells using the periodate oxidation and aniline-catalyzed oxime ligation (PAL) method. (c) Flow cytometry histograms and quantification of diCBM40 levels on MCF7. (d,e) Flow cytometry histograms and quantification of SLBR-N levels on MCF7 (d) and BoM-1833 (e); SLBR-N levels for MCF7 quantified by measurement of mean fluorescent intensity in confocal microscopy images. (f,g) Flow cytometry analysis of Hyaluronic Acid Binding Protein (HABP) level on MDA-MB231 (f) and MCF7 (g). HABP levels quantified by measurement of mean fluorescent intensity in confocal microscopy images. In panel **a**, **d**, **f**, and **g**, results are the mean ± s.d. ($n$ = 3) and statistical differences were determined using two-tailed unpaired t-test.

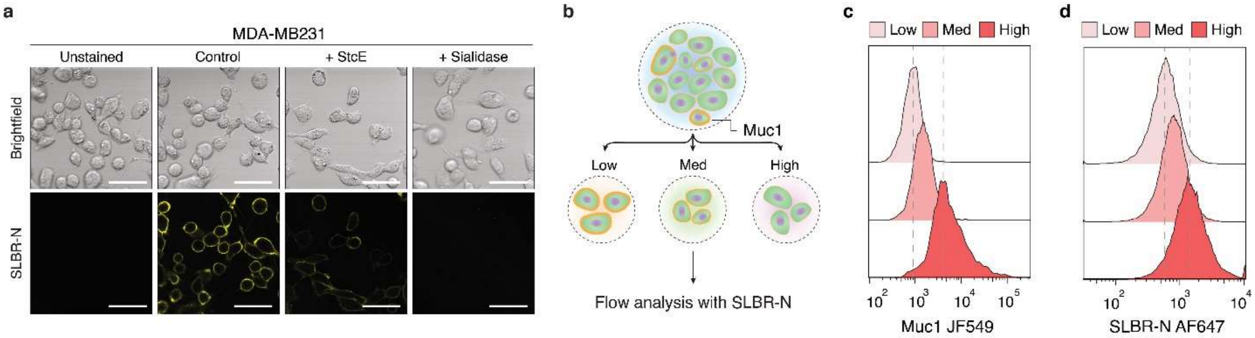


**Supplementary Figure 4. SLBR-N binds sialylated mucin O-glycans.** (a) Representative brightfield and confocal images of MDA-MB231 cells labelled with SLBR-N conjugated with Alexa Fluor 647. MDA-MB231 cells were treated with control buffer or 100 nM StcE or 100 nM sialidase in cell culture medium for 1 hour. (b) Schematic showing the MDA-MB231 cells were sorted into three sub-populations with low, intermediate, and high levels of endogenous Muc1 expression. The cells were then labelled with SLBR-N conjugated with Alexa Fluor 647. (c,d) Flow cytometry analysis of Muc1 (c) and SLBR-N level (d) on MDA-MB231 cells after sorting.

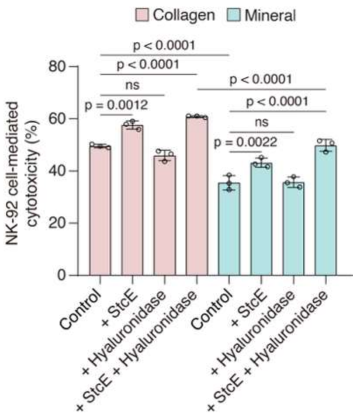


**Supplementary Figure 5. Cell-surface hyaluronan does not mediate protection against effector cells.** NK-92 cell-mediated cytotoxicity of MDA-MB231 treated with control buffer, 2.5 U/mL *Streptomyces hyalurolyticus* hyaluronidase or 100 nM StcE mucinase in cell culture medium for 1 hour prior to co-culture. E:T ratio is 10:1. Results are the mean ± s.d. ($n = 3$). Statistical differences were determined using one-way ANOVA with multiplicity-adjusted p values from Tukey's multiple comparisons test.
